# The dynamic transmission of positional information in *stau^-^* mutants during *Drosophila* embryogenesis

**DOI:** 10.1101/868711

**Authors:** Zhe Yang, Hongcun Zhu, KaKit Kong, Jiayi Chen, Xiaxuan Wu, Peiyao Li, Jialong Jiang, Jingchao Zhao, Feng Liu

**Affiliations:** State Key Laboratory of Nuclear Physics and Technology & Center for Quantitative Biology, Peking University, 100871 Beijing, P. R. China.

## Abstract

Intriguingly, the developmental patterning during *Drosophila* embryogenesis is highly accurate and robust despite its dynamic changes and constant fluctuations. It has been suggested that Staufen (Stau) is key in controlling the boundary variability of the gap protein Hunchback (Hb). However, its underlying mechanism is still elusive. Here, we have developed methods to quantify the dynamic 3D expression of segmentation genes in *Drosophila* embryos. With improved control of measurement errors, our results reveal that the posterior boundary of the Hb anterior domain (*x_Hb_*) of *stau^-^* mutants shows comparable variability to that of the wild type (WT) and shifts posteriorly by nearly 12% of the embryo length (EL) to the WT position in the nuclear cycle (nc) 14. This observed large shift might contribute significantly to the apparent large variability of *x_Hb_* in previous studies. Moreover, for *stau^-^* mutants, the upstream Bicoid (Bcd) gradients show equivalent gradient noise to that of the WT in nc12-nc14, and the downstream Even-skipped (Eve) and cephalic furrow (CF) show the same positional errors as the WT. Our results indicate that threshold-dependent activation and self-organized filtering are not mutually exclusive but could both be implemented in early *Drosophila* embryogenesis.

## Introduction

During the development of multicellular systems, the expression of patterning genes dynamically evolves and stochastically fluctuates (Dubuis et al., 2013; Gregor et al., 2007b; Jaeger et al., 2004; Kanodia et al., 2009; Liu et al., 2013; Yang et al., 2018). It is intriguing how developmental patterning achieves a high degree of accuracy (Dubuis et al., 2013; Gregor et al., 2007a) and robustness (Inomata et al., 2013; Liu et al., 2013; Lucchetta et al., 2005). Two hypotheses have been proposed: one is the threshold-dependent model, i.e., the French flag model, which assumes that the positional information is faithfully transferred from precise upstream patterning (Rogers and Schier, 2011; Wolpert, 2011); the other is the self-organized filtering model, which assumes that noisy upstream patterning needs to be refined to form downstream patterning with sufficient positional information (Dubuis et al., 2013; Gregor et al., 2007b; Houchmandzadeh et al., 2002; Jaeger et al., 2004; Kanodia et al., 2009; Liu et al., 2013; Manu et al., 2009; Yang et al., 2018). The two models have been thought to be mutually exclusive, and it has been extensively debated which one is implemented in a particular developmental system.

The *Drosophila* embryo is an excellent model system to address this question. It is believed that the blueprint of the adult body plan is established during the first 3-hour patterning before gastrulation in embryos. In particular, the adult body segments can be mapped with the expression pattern of the segmentation genes along the anterior and posterior (AP) axis. The hierarchic segmentation gene network consists of four layers of patterning genes (Gregor et al., 2007a; Jaeger, 2011): maternal morphogen such as *bicoid* (*bcd*) (Dubuis et al., 2013; Porcher and Dostatni, 2010; Struhl et al., 1989), zygotic gap genes such as *hunchback* (*hb*) (Inomata et al., 2013; Liu et al., 2013; Lucchetta et al., 2005; Struhl et al., 1992), pair rule genes such as *even-skipped* (*eve*) (Goto et al., 1989; Rogers and Schier, 2011; Wolpert, 2011), and segmentation polarity genes (Swantek, 2004). They form increasingly refined developmental patterns along the AP axis until the positional information carried by the patterning genes reaches the single-cell level, i.e., 1% embryo length (EL), in the variability of the expression pattern (Dubuis et al., 2013; Gregor et al., 2007a).

It has been controversial which hypothesis is valid in the dynamic transmission of positional information during *Drosophila* embryogenesis, and the Hb boundary has been an important subject in this investigation (Erclik et al., 2017; Gregor et al., 2007a; Houchmandzadeh et al., 2002; Huang et al., 2017; Lucas et al., 2018; Staller et al., 2015; Tran et al., 2018). Lying directly downstream of maternal gradients, zygotic Hb forms a steep boundary at approximately the middle of the embryo with a variability of 1% EL. The variability of the position of the Hb boundary (*x_Hb_*) has long been thought to depend on the gene *staufan* (*stau*) (Houchmandzadeh et al., 2002). As shown in previous work, *stau* was the only gene that dramatically increased the variability of *x_Hb_* from 1% EL to more than 6% EL for protein profiles (Houchmandzadeh et al., 2002) and 4% EL for mRNA profiles (Crauk and Dostatni, 2005). In contrast, the variability of *x_Hb_* remains almost the same as that of the wild type (WT) with knockout of nearly all the genes that potentially interact with Hb, including *nos*, *Kr*, and *kni*, or even deletion of the whole or half of the chromosome (Houchmandzadeh et al., 2002). Hence, *stau* could be the key to understanding the potential noise-filtering mechanism.

However, the underlying mechanism of the role of *stau* has been elusive. It is well known that Stau does not interact with Hb directly but is necessary for the localization of the *bcd* mRNA at the anterior pole (Ferrandon et al., 1994) and the spatially constrained translation of *nos* mRNA at the posterior pole (St Johnston et al., 1991). Hence, Stau is important for establishing the Bcd gradient that activates the transcription of Hb (Struhl et al., 1989) and the Nos gradient that represses the translation of maternal Hb (Wang and Lehmann, 1991). The role of Bcd in regulating the variability of the Hb boundary remains controversial. On the one hand, Houchmandzadeh et al. suggested that the variability of *x_Hb_* was independent of upstream Bcd gradients, as the average Bcd gradients of two groups of embryos overlapped, although their Hb shifted anteriorly or posteriorly compared with the average Hb profile, and the variability of *x_Hb_* was much smaller than the positional error derived from the Bcd gradient noise (Houchmandzadeh et al., 2002). On the other hand, He and et al. showed that the shift of *x_Hb_*correlated with the average Bcd gradient showing different concentrations at *x_Hb_*, and the variability of *x_Hb_* was equivalent to the positional error derived from the Bcd gradient noise (He et al., 2008). In addition, the variability of *x_Hb_*seemed to be significantly less in He’s measurement (He et al., 2008) than in Houchmandzadeh’s measurement (Houchmandzadeh et al., 2002).

Some of these controversial points might be clarified if we could further reduce the spatial and temporal measurement errors in developmental patterning. It has been estimated that the orientation-related error could be as high as 20-50% of the total measurement error for Bcd gradients (Gregor et al., 2007a) and gap genes (Dubuis et al., 2013). This is because the dynamic developmental profiles vary spatially in the asymmetric 3D embryos, yet traditionally they are usually measured in a selected plane of the manually oriented embryo (Gregor et al., 2007a; Houchmandzadeh et al., 2002).

Here, we developed measurement methods to quantify the dynamic 3D expression of patterning genes in *Drosophila* embryos and applied these methods to measure the positional errors of the segmentation genes at different levels. We chose one of the strongest *stau* alleles, *stau^HL54^*, which induces the largest variability of *x_Hb_* (Houchmandzadeh et al., 2002), as the *stau^-^* mutant in this study. Surprisingly, we discovered that in *stau^-^* mutants, *x_Hb_*shifts posteriorly by nearly 12% EL to the WT position in nc14, and the variability of *x_Hb_* in an ∼5-min time interval is comparable with that of the WT most of the time in nc14. Moreover, the upstream Bcd gradients show equivalent gradient noise to that of the WT from nc12-nc14, and the downstream Eve and CF show the same positional errors as the WT. We also constructed a minimal model and revealed that the extremely large shift in the Hb boundary in *stau^-^* mutants originates from the Bcd gradients with a 50% decrease in amplitude and a 17% increase in length constant in *stau^-^* mutants compared with the WT. Our results indicate that the threshold-dependent model could be valid at early nc14, but then the gene network adjusts, and the filtering mechanism is implemented at least from the maternal Bcd gradient to Hb. Some factors other than *stau,* which remain to be discovered, could play important roles in filtering dynamic positional information.

## Results

### A large dynamic shift in the Hb boundary of *stau^-^*mutants

To reduce the measurement errors, we quantified the Hb expression pattern using 3D imaging with light-sheet microscopy (Figure 1-figure supplement 1, Figure 1-video 1, Figure 1-video 2) on fixed and immunostained embryos and staged each embryo with 1-min temporal precision using the depth of the furrow canal (Figure 1-figure supplement 2). The average temporal Hb expression profiles of the WT (Figure 1-figure supplement 3) and *stau^-^* mutants (Figure 1A) are shown on heat maps projected from 3D embryo images. From the heat map, we extracted the average of the normalized dorsal profile of Hb with a time-step of 5 min in nc14 (Figures 1B-C) and measured *x_Hb_*. Consistent with previous results, *x_Hb_*of the WT remains nearly constant after 20 min into nc14. Notably, in the first 20 min into nc14, it shifts posteriorly by 3% EL (Figure 1D). In contrast, at the dorsal side of the embryo, *x_Hb_* of *stau ^-^* mutants starts at 35.9 ± 1.1% EL at 5 min into nc14, then dynamically shifts posteriorly by 11.3% EL within 60 minutes, and finally stabilizes at 47.2 ± 1.1% EL, close to the WT boundary position 48.4 ± 0.8% EL (Figure 1D, Figure 1-supplement 4C and E). Moreover, *x_Hb_*differs significantly in different orientations, e.g., the ventral boundary moves from 29.7 ± 1.2% EL by 11.6% EL to 41.3 ± 1.2% EL, shifting anteriorly by approximately 6% EL compared with the dorsal boundary (Figure 1D, Figure 1-figure supplement 4A, D, and F). These results suggest that it is crucial to control spatial and temporal measurement errors to assess the variability of the Hb boundaries. Interestingly, the spatial measurement error of *x_Hb_* seems to be the smallest on the dorsal profiles (Figure 1-figure supplement 5A-B).

**Figure 1.**
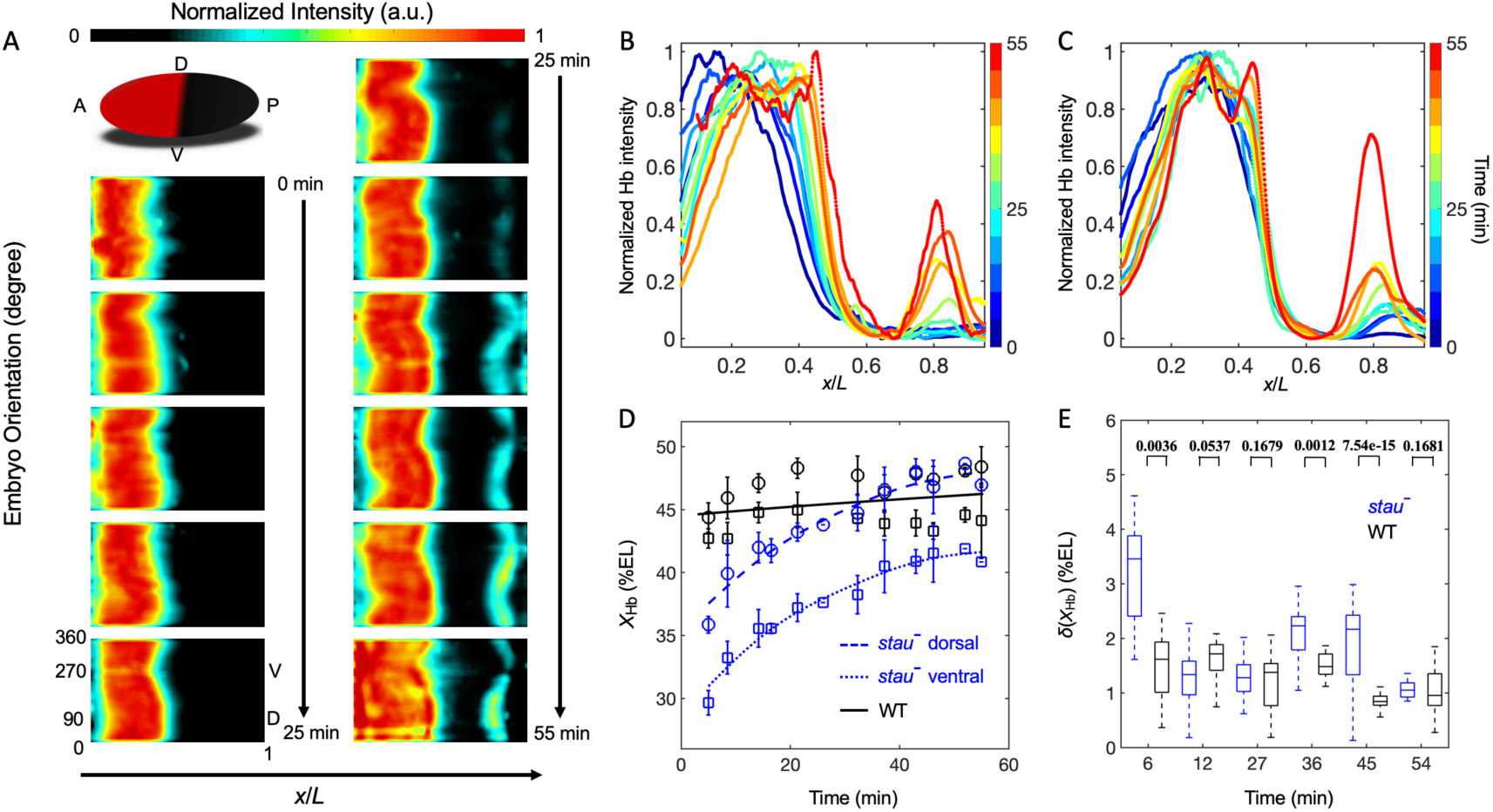
Bcd gradient noise of *stau^-^* mutants is comparable with that of the WT. (A) Heat map of the normalized Hb intensity in *stau^-^*mutants with a time-step of 5 min in nc14 based on projection at different angles along the AP axis from the 3D imaging on embryos. (B-C) Dynamics of the average dorsal Hb profiles extracted from 3D imaging on the embryos of *stau^-^*mutants (B) and the WT (C) in nc14. (D) The average Hb boundary position (*x_Hb_*) in each 5-min bin of *stau^-^* mutants (blue) and the WT (black) as a function of the developmental time in nc14. Each circle (square) represents the average of the dorsal (ventral) boundary positions of Hb in the bin, and error bars denote the standard deviation of the bin. Lines are eye guides. (E) The variability of the Hb boundary in each bin with equal sample numbers (*N*=6) of *stau^-^*mutants (blue) and the WT (black) as a function of the developmental time in nc14. Error bars are calculated from bootstrapping. The *p* values of the F-test of the statistical significance of the difference between the Hb boundary variability of *stau^-^* mutants and the WT are shown at the top.

In fact, in ∼5 min time windows in nc14, the variability of the Hb boundary of *stau^-^* mutants is less than 2.5% EL except in very early nc14. This difference was not statistically significant from the WT from 12 min to 36 min into nc14 (Figure 1E). This result is different from previous measurements. We suspect that the apparent difference in variability of *x_Hb_* might result from the different control of the measurement errors (Figure 1-figure supplement 5C-D), e.g., the temporal control in He’s measurements (He et al., 2008) could be better than Houchmandzadeh’s measurements (Houchmandzadeh et al., 2002), and the spatial control could be better in our measurements than in He’s measurements (He et al., 2008).

### Bcd gradient noise of *stau*^-^ mutants remains nearly the same as that of the WT

To understand the origin of the variability of the Hb boundary, we measured its upstream Bcd gradients. Using live imaging with a two-photon microscope (TPM), we found that the average Bcd-GFP gradient of *stau^-^* mutants at 16 min into nc14 shows reduced amplitude but an increased length constant (Figure 2A). Compared with the WT, the amplitude of the Bcd-GFP gradient in *stau^-^*mutants is only 48±7%, but the length constant is increased by approximately 13% from 18.6% EL to 21.1% EL (Figure 2A). Notably, after correcting for the GFP maturation effect (Little et al., 2011; Liu et al., 2013), the length constant of the Bcd gradient of *stau^-^*mutants increases by approximately 17% from 15.5% EL of the WT to 18.2% EL (Figure 2-figure supplement 1B). This is reasonable because the *bcd* mRNA is more extensively distributed in the embryos of *stau^-^* mutants (Ferrandon et al., 1994; Petkova et al., 2014). Interestingly, the Bcd gradient noise of *stau^-^* mutants is comparable with that of the WT (Figure 2B). The relative noise of the Bcd-GFP gradient is 15.0±3.6%, which is very close to 15.0±5.4% of the WT. Hence, it seems to be consistent with the comparable variability of the Hb boundary. However, this result is different from the previous measurement (He et al., 2008; Houchmandzadeh et al., 2002), which might have different measurement errors.

**Figure 2.**
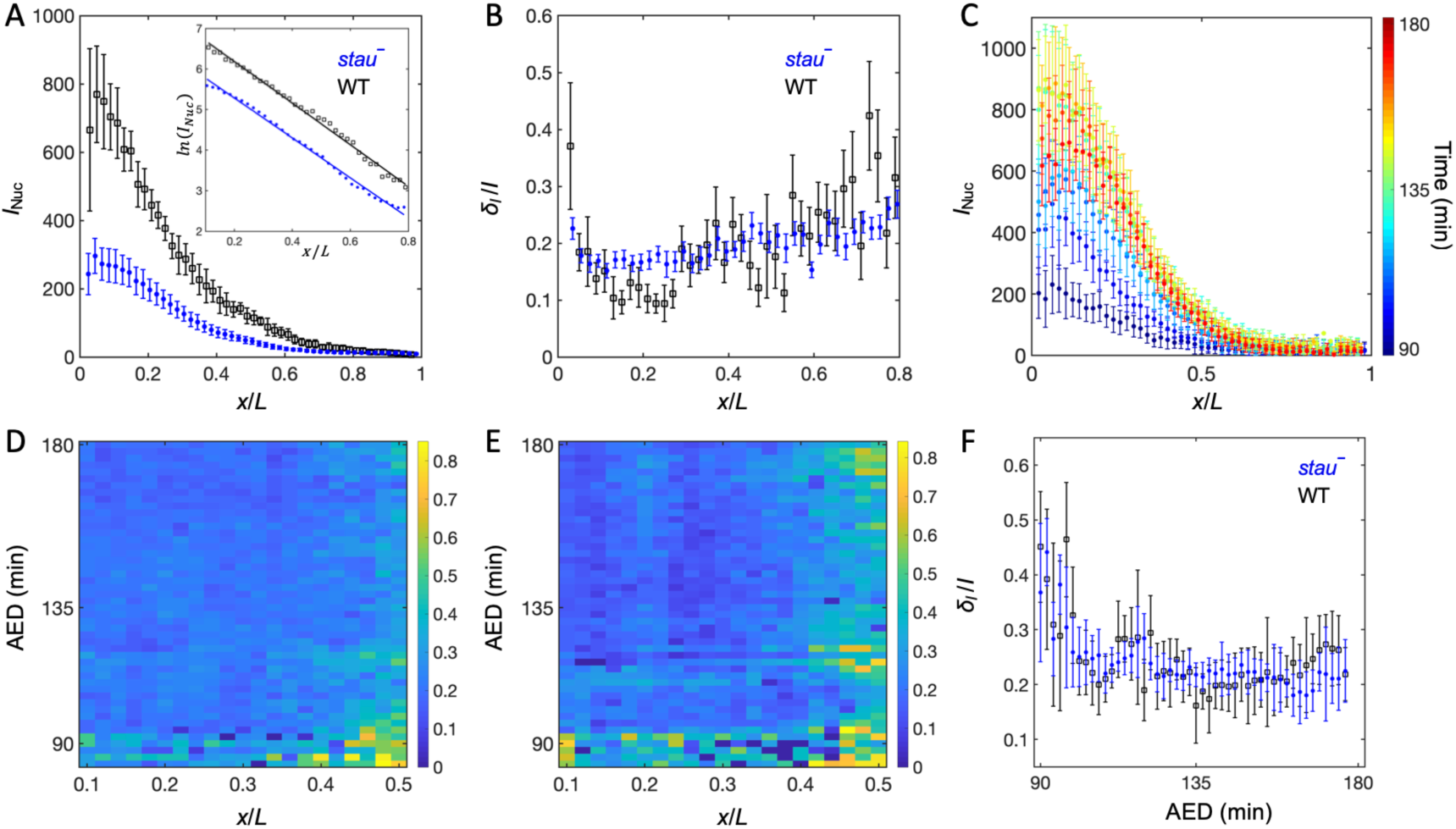
The Bcd gradient noise of *stau^-^* mutants is comparable with that of the WT. (A) The average intensity of the nuclear Bcd-GFP gradient of *stau^-^* mutants (blue) and the WT (black) as a function of the fractional embryo length at 16 min into nc14. Each circle and error bar represent the average and standard deviation of the Bcd-GFP fluorescence intensity of the nuclei in the bin with a bin size of 2% EL, respectively. Inset shows the logarithm of the intensity as a function of the fractional embryo length and the linear fit. (B) The relative Bcd-GFP gradient noise of *stau^-^* mutants (blue) and the WT (black). Each circle represents the standard deviation divided by the mean of the Bcd-GFP fluorescence intensity of the nuclei in the bin with the bin size of 2% EL. The error bar represents the standard deviation of the mean relative gradient noise calculated with bootstrap. (C) Variation in the average profiles of the Bcd gradient in *stau^-^* mutants from nc12 to 60 min into nc14. (D-E) The heat map of the Bcd gradient noise of the WT (D) and *stau^-^*mutants (E) from nc12 to 60 min into nc14. (F) The average Bcd gradient noise of *stau^-^* mutants from *x*=0 to *x*=0.6 vs. developmental time is comparable with that of the WT.

Another concern is that Bcd-dependent Hb expression turns off in less than 10 min into nc14 (Liu and Ma, 2013; Liu et al., 2016), and the time window for Bcd interpretation could be earlier than nc14 (Huang et al., 2017; (Bergmann et al., 2007), hence Hb expression could be largely obtained by translating the previously produced mRNA. It is therefore necessary to measure the dynamic Bcd gradient noise in developmental time earlier than the conventional measurement time, i.e., 16 min into nc14. By improving the imaging technique and image analysis method (Figure 2-figure supplement 1), we assessed the Bcd-GFP gradient noise in both *stau^-^* mutants and the WT. The average nuclear Bcd-GFP gradient rises to a maximum in nc14 and then slightly falls off (Figure 2C). This is consistent with the results of another live imaging experiment (Gregor et al., 2007b) but is significantly different from the measurement of the fixed embryos (Little et al., 2011) due to GFP maturation effect (Little et al., 2011; Liu et al., 2013). Importantly, the gradient noise of *stau^-^* mutants and the WT remains comparable from nc12 to 60 min into nc14 (Figure 2D-F). It is also interesting to discover that the Bcd-GFP gradient noise in both fly lines remains nearly the same level from nc13 to nc14 (Figure 2F). These conclusions should still be valid even if we consider the increased measurement errors (probably due to imperfect fluorescence intensity calibration in different sessions), which slightly increase the gradient noise at 16 min into nc14 compared with the one with TPM (Figure 2B), as the measurement error should change at the same level for both *stau^-^* mutants and the WT.

### The threshold-dependent activation model is valid in early nc14

Similar to *stau^-^* mutants, the amplitude of the Bcd-GFP gradient in the fly line with only half of the Bcd dosage (Bcd1.0) also decreases by half compared with the WT (Liu et al., 2013). According to a previous study, for the fly line Bcd1.0, the Hb boundary position might be determined according to the threshold-dependent activation model in early nc14. Then, it dynamically shifts posteriorly toward the Hb position of the WT but stops midway in later nc14. In contrast, the Hb position in *stau^-^* mutants shifts all the way to the WT position (Liu et al., 2013). It is interesting to investigate whether the threshold-dependent activation model is also valid in early nc14 in *stau^-^* mutants.

To test this speculation, we first needed to correct the GFP maturation effect to obtain the total Bcd-GFP gradient from the observed Bcd-GFP gradient measured with live imaging, as it takes tens of minutes for the newly synthesized Bcd-GFP to fluoresce. Based on the synthesis-degradation-diffusion (SDD) model incorporating maturation correction and the spatial distribution of *bcd* mRNA (Little et al., 2011; Petkova et al., 2014), we calculated the maturation correction curves for the Bcd-GFP gradient in both *stau^-^* mutants and the WT (Figure 3-figure supplement 1A). Interestingly, the Bcd-GFP concentrations at the respective Hb boundary position are almost the same at 16 min into nc14 (Figure 3), suggesting that the threshold-dependent model might still apply in early nc14. However, as *x_hb_* of *stau^-^* mutants shifts posteriorly in a much greater range than the WT, and the Bcd gradient changes only slightly in both fly lines, it remains to be tested in which exact time window this model applies.

**Figure 3.**
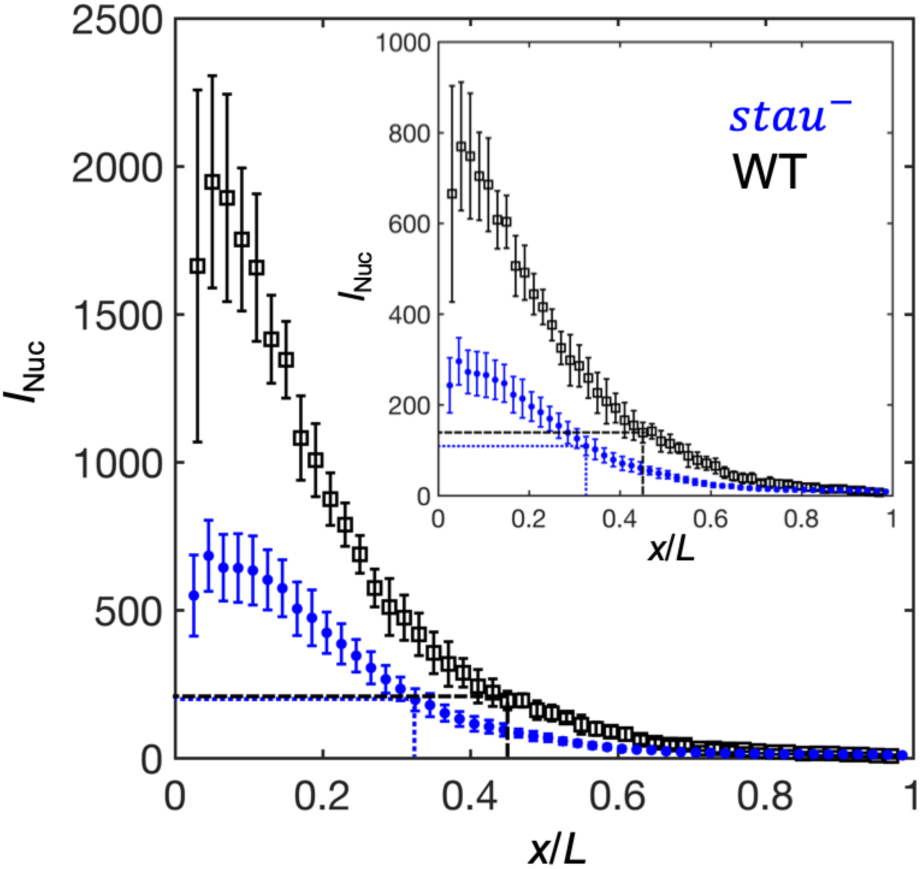
The activation of Hb by Bcd is consistent with the threshold-dependent model in *stau^-^* mutants at early nc14. Comparison of the Bcd concentration (horizontal lines) at *x_Hb_* (vertical lines) at 16 min into nc14 between *stau^-^* mutants (blue) and the WT (black) before (inset) and after maturation correction on the Bcd gradient. Each circle and error bar represent the average and standard deviation of the Bcd-GFP fluorescence intensity of the nuclei in the bin with a bin size of 2% EL, respectively.

### Positional information transmission

In addition to the threshold-dependent model, it is also interesting to test whether the noise-filtering model plays a role in early *Drosophila* embryogenesis. We measured the positional information transmission of the other patterning genes interacting with Hb in *stau^-^*mutants. As another gap protein, Kr forms a strip adjacent to the Hb boundary, and the two genes suppress each other (Jaeger, 2011). In *stau^-^*mutants, the Kr strip narrows (Figure 4-figure supplement 1A) and shifts posteriorly together with the Hb boundary in early nc14 (Figure 4A) but stops movement late, suggesting that the late shift in the Hb boundary is independent of Kr. As one of the downstream pair rule proteins, Eve forms only 4 instead of 7 strips in *stau^-^* mutants (Figure 4-figure supplement 2B). The dynamic shift in the four strips is also shown in Figure 4A. At 56 min into nc14, the cephalic furrow (CF) position in the *stau^-^*mutants is 28.2±1.1% EL, while the CF of the WT is 34.3±1.2% EL.

**Figure 4.**
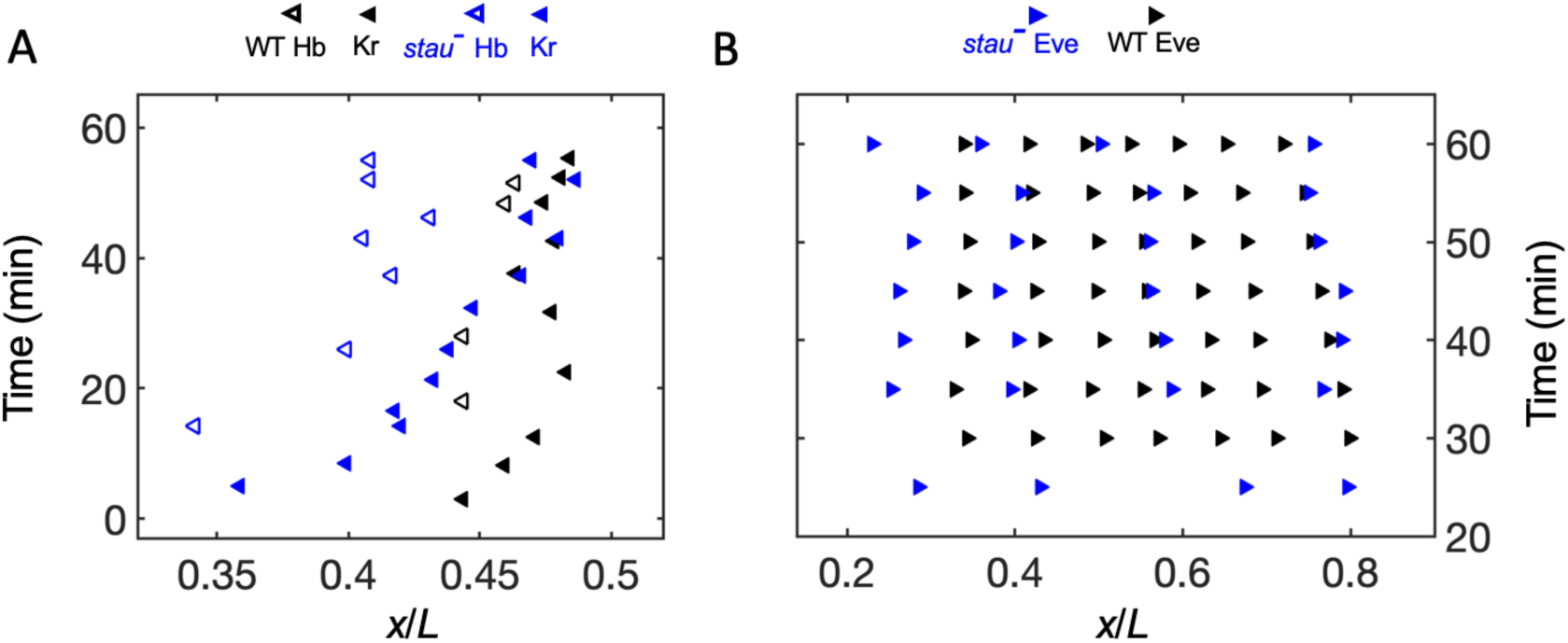
The position variation in Kr and Eve is much smaller than that in Hb in *stau^-^* mutants. (A) The positions of the anterior boundary (inflection point) of Kr (solid triangle) and *x_Hb_* (hollow triangle) with the *stau^-^* mutants (blue) and the WT (black). (B) The peak of Eve in *stau^-^* mutants (blue) as a function of the developmental time in nc14 in comparison with the WT (black).

To understand the dynamic positional information transmission, we compared the positional noise of all the measured AP patterning markers from Bcd to CF during the time course of nc14 (Figure 4C, Figure 4-figure supplement 2). Consistent with previous results, the positional error of the Hb boundary of the WT decreases by approximately 2-fold to a minimum at approximately 45 min into nc14 and then slightly increases (Dubuis et al., 2013). The *stau^-^* mutant appears to follow a slightly different trend: it shows a positional error of 3.5% EL at 6 min into nc14, reaches a minimum of 1.2% EL at approximately 20 min into nc14, rises back to approximately 2% EL at 45 min into nc14, and finally decreases to approximately 1% EL at 56 min into nc14. The positional error of *x_Hb_*in early nc14 is comparable with that of the Bcd gradient in the WT, but slightly greater in *stau^-^* mutants. On the other hand, the minimal positional errors of the Kr peak, the first peak of Eve, and CF are comparable with the minimal positional error of the Hb. Notably, the *stau^-^*mutants and the WT are comparable in their minimal positional errors in different levels of AP patterning genes (Table S1). These results suggest that positional noise is filtered from Bcd to Hb and then relayed between different levels of the patterning genes.

In addition to the noise-filtering phenomenon, it is interesting to note that the shifts in the patterning markers differ. The Hb boundary shifts posteriorly by nearly 12% EL to the WT position. For the other gap genes that interact with Hb, the front boundary of Kr shifts posteriorly by only 5% EL and stops 3% EL away from the WT position. For the downstream genes, the first peak of Eve shifts very little, and its final position corresponding to the CF position is 6% EL away from the WT position. These results indicate that the large dynamic shift of Hb seems to be dampened instead of faithfully relayed in the other patterning genes, in contrast to the shift accumulation in the fly lines with altered Bcd dosages (Liu et al., 2013). For example, in the fly line Bcd1.0 with almost the same Bcd dosage as *stau^-^*mutants, the Hb boundary position shifts posteriorly by approximately 2.5% EL in nc14 and stops at 7% EL away from the WT position. The first peak of Eve shifts posteriorly further by 0.8% EL, so the CF position is closer to the WT position. It has been suggested that this additive shift results from the dynamic integration of maternal positional information. It remains to be investigated whether a new mechanism may account for the dynamic positional transmission in *stau^-^* mutants.

## Discussion

### Improving quantitative measurement methods to reveal the true biological noise of developmental patterning

Maternal mutants become more accessible with the CRISPR-Cas9 technique (Bassett and Liu, 2014). The balancer stabilizes the maternal mutant fly line but prevents the random mutation from being rescued by homologous recombinations. Hence, the accumulated mutations often degrade the maternal mutant fly line in the fly stock. In this work, instead of running rescue genetics experiments, we generated a fresh *stau^-^* mutant fly line from the WT with the CRISPR-Cas9 technique. The newly generated *stau^HL54^*mutant fly line shows the correct cuticle (data not shown) and expression patterns of Bcd, Hb, Kr, and Eve compared with the original *stau^-^*mutant fly line (Figure 4-figure supplement 2).

To reveal the true biological noise, 3D imaging is very helpful in controlling spatial measurement errors. Compared with the two-photon microscopy used in previous studies for 3D imaging on embryos (Fowlkes et al., 2008), the light-sheet microscope should provide better-quality 3D reconstruction by combining images taken from two opposite directions with higher imaging speed (Krzic et al., 2012). With the 3D imaging data, we can evaluate the measurement error in embryo orientations. Interestingly, most of the time, the measurement errors at the dorsal side are smaller than the ventral side and the symmetric two sides on the coronal plane (Figure 1-figure supplement 5A-B). This is because the Hb boundary shifts in a much smaller range around the dorsal side (Figure 1A, Figure 1-figure supplement 3). Nevertheless, even with 10° uncertainty in the orientation, the measurement error at the dorsal side could still be as high as 0.2% EL, at least 20% of the positional error of the Hb boundary, which is 1% EL.

For dynamically evolved patterning, it is also important to precisely determine the embryo age. Both the depth of the furrow cannel (FC) and the nuclear shape/size are good measures for the embryo age in nc14. The former (latter) has better time resolution in the late (early) developmental stage. Notably, the dynamic shift in FC could vary in different fly lines, as the shift curve measured with *w^1118^* is different from previously published results measured with Oregon-R (Figure 1-figure supplement 2A)(Dubuis et al., 2013).

Compared with imaging fixed embryos, live imaging has advantages in higher temporal resolution but is often prone to slow imaging speed. Traditionally, the Bcd gradient noise is often measured only at 16 min into nc14 in a selected plane, as this is the most convenient in live imaging. However, the Bcd-dependent regulation of Hb has already been shut off before the Bcd gradient is measured (Liu and Ma, 2013). Moreover, considering the dynamics of downstream patterning genes, it is important to measure the dynamics of the Bcd gradient noise. However, before nc14, additional fluorescence intensity measurement errors could be introduced because some of the sparsely distributed nuclei on the embryo surface may be offset from the imaging plane. We found that the maximum projection from a 5-layer and 1 µm-spaced z-stack image could significantly alleviate the intensity of measurement errors. To accumulate sufficient samples to measure the gradient noise, it is also very important to stabilize the imaging condition and correct the potential intensity drift between different experimental sessions (Figure 2-figure supplement 1C-D). With this imaging improvement as well as automatic image analysis, we successfully showed that the Bcd-GFP gradient noise is almost constant in the interphase from nc13 to nc14 (Figure 2F). Since the fluorescence intensity observed in live imaging increases in this period, this result may suggest that the Bcd gradient noise is not dominated by the Poissonian noise.

### Underlying mechanism for the dynamic shift of the Hb boundary

Based on the quantitative spatial-temporal gene expression data, it has long been known that the gap gene profiles, e.g., central Kr domain and the posterior Kni and Gt domain, show substantial anterior shifts during nc14 (Jaeger et al., 2004). These dynamic shifts are proposed to originate from the asymmetric cross-regulation between gap genes, i.e., posterior gap genes repress their adjacent anterior gap genes but not vice verse (Jaeger et al., 2004). The shift amount is less than 5%, much less than the shift of the Hb boundary in *stau^-^* mutants.

The Hb boundary and the other patterning features, such as the Kr central strip and Eve peaks, also shift dynamically with differing Bcd dosages (Liu et al., 2013). For example, for the fly line with only half of the Bcd dosage in the WT, the shift in the CF is only approximately 40% of the predicted value based on the threshold-dependent model. The Hb boundary shifts posteriorly toward the WT position by approximately 4% EL from an initial position close to the one predicted by the threshold-dependent model. The posterior shift of the Hb boundary has also been observed in *nos^-^* mutants (Petkova et al., 2019).

Among all the reported patterning shift dynamics in fly embryos, the shift in the Hb boundary in *stau^-^* mutants is the largest at nearly 12% EL. The shift significantly slows down in late nc14, as the majority of the shift, ∼ 9% EL, finishes in the first half an hour of nc14. In contrast, most of the shifts observed in the other cases occur in the later half an hour in nc14 and have been proposed to originate from the cross-regulation between gap genes (Jaeger et al., 2004; Liu et al., 2013). Hence, a mechanism other than the cross-regulation between gap genes could contribute to the large shift of the Hb boundary in *stau^-^* mutants. Moreover, the slope of the Hb boundary in *stau^-^* mutants is less than that of the WT in the first half an hour in nc14 and later increases to nearly the same as that of the WT (Figure 5-figure supplement 1). This might be related to the modification of the maternal gradients from both poles (Figs. 6 A-B). On the one hand, the amplitude of the Bcd gradient is reduced by half, and the length constant is increased by 17%. On the other hand, the maternal Hb profile is flattened as the Nos gradient is removed.

To test this idea, we constructed a minimal mathematical model to calculate the dynamic shift in *x_Hb_*, taking into account both the activation from Bcd to Hb and the self-activation of Hb (for more details, see Supplementary Materials and Figure 6-figure supplement 1). This model fits well with the measured data (Figure 6C). Based on this model, the shift in *x_Hb_* from the initial position to the final position in nc14 follows an exponential trend, and the time constant of *stau*^-^ mutants is approximately twice that of the WT. This ratio is consistent with the model prediction if we assume the activation of Hb by Bcd follows the Hill function with a Hill coefficient of 5 (Gregor et al., 2007a) and use the maturation corrected Bcd gradients as the input.

### Dynamic positional information transmission

The large dynamic shift in the Hb boundary raises the question of how the positional information is transferred in the patterning system. The term “positional information” was first coined by Wolpert (Wolpert, 1969). Based on this concept, developmental patterning is instructed by the concentration of a single static morphogen gradient, and the interpretation of the morphogen gradient follows the threshold-dependent model, i.e., each cell “acquires” its position inside the embryo through “reading” the morphogen concentration and accordingly activates downstream genes to form the cell fate map (Jaeger and Reinitz, 2006; Wolpert, 2011; 2016). This model provides a simple molecular-based mechanism for developmental pattern formation. It has prevailed in developmental biology, especially after the identification of a series of morphogens starting with Bcd (Rogers and Schier, 2011; Struhl et al., 1989).

However, this model has long been challenged because gradient noise could disrupt patterning precision (Houchmandzadeh et al., 2002; Jaeger et al., 2007). Without precise morphogen gradients as inputs, the developmental pattern could also form via a noise-filtering mechanism resulting from cross-regulation between genes (Manu et al., 2009). This idea is rooted in Turing’s seminal idea: periodic patterns can spontaneously form in a self-organizing reaction-diffusion system, e.g., a slow diffusive activator and a fast diffusive inhibitor (Corson and Siggia, 2012; Turing, 1952). Recently, an increasing number of developmental patterning systems have been found to implement a Turing-like mechanism (Economou et al., 2012; Goryachev and Pokhilko, 2008; Raspopovic et al., 2014).

Usually, these two classic mechanisms have been thought to be mutually exclusive. Hence, great effort has been made to distinguish which mechanism is implemented in a particular developmental system. However, it has been controversial whether early fly embryogenesis follows the threshold-dependent model or the noise-filtering model (Gregor et al., 2007a; He et al., 2008; Houchmandzadeh et al., 2002; Manu et al., 2009) because it has been challenging to run quantitative tests. First, it is well known that the gap gene dynamically changes in nc14 (Dubuis et al., 2013; Jaeger et al., 2004). Hence, the developmental system is not in steady state, and the concentration of these transcription factors could keep changing with a time scale much shorter than the degradation time of the downstream gene. As a result, the developmental pattern cannot be regarded as the instant readout of simultaneous transcription factors but as an accumulation with the product generated in an early time window. Hence, it would be misleading to test the threshold-dependent model based only on the snapshot of the protein concentration profiles. For a strict test of the Bcd-Hb activation system, ideally, we need to measure the profile of the *hb* transcription activity and the Bcd gradient at an even earlier time before the Bcd-dependent transcription turns off. Second, the gap gene integrates multiple maternal factors (Jaeger, 2011; Liu et al., 2013). Hence, we need to measure the combined positional information of all the upstream genes and the dynamic positional information of the downstream genes. However, the regulatory function is still unknown, although the combined positional information might be estimated based on the optimal decoding hypothesis (Petkova et al., 2019). Moreover, the conventional measurement method without sufficient spatial and temporal measurement error control is rather limited in measuring the expression dynamics.

By developing 3D measurements with reduced measurement errors, we observe that the positional errors decrease from approximately 3.5% EL at early nc14 to approximately 1% EL in the middle of nc14 in *stau^-^* mutants. A slight decrease in the positional errors was also observed in the WT (Figure 5). Interestingly, even for *stau^-^* mutants with an ultra-large dynamic shift in the Hb boundary, the threshold for Bcd in activating Hb is still the same as that of the WT at 16 min into nc14. These results suggest that the threshold-dependent model probably acts upstream of the filtering mechanism during *Drosophila* embryogenesis. It has also been suggested that Wolpert’s positional information model, i.e., the threshold-dependent morphogen model, and the self-organized Turing model could coexist in other developmental patterning systems (Green and Sharpe, 2015). We expect that studying the combination of the two mechanisms could be the key to revealing the mechanism of precise and robust pattern formation via dynamic positional information transmission.

**Figure 5.**
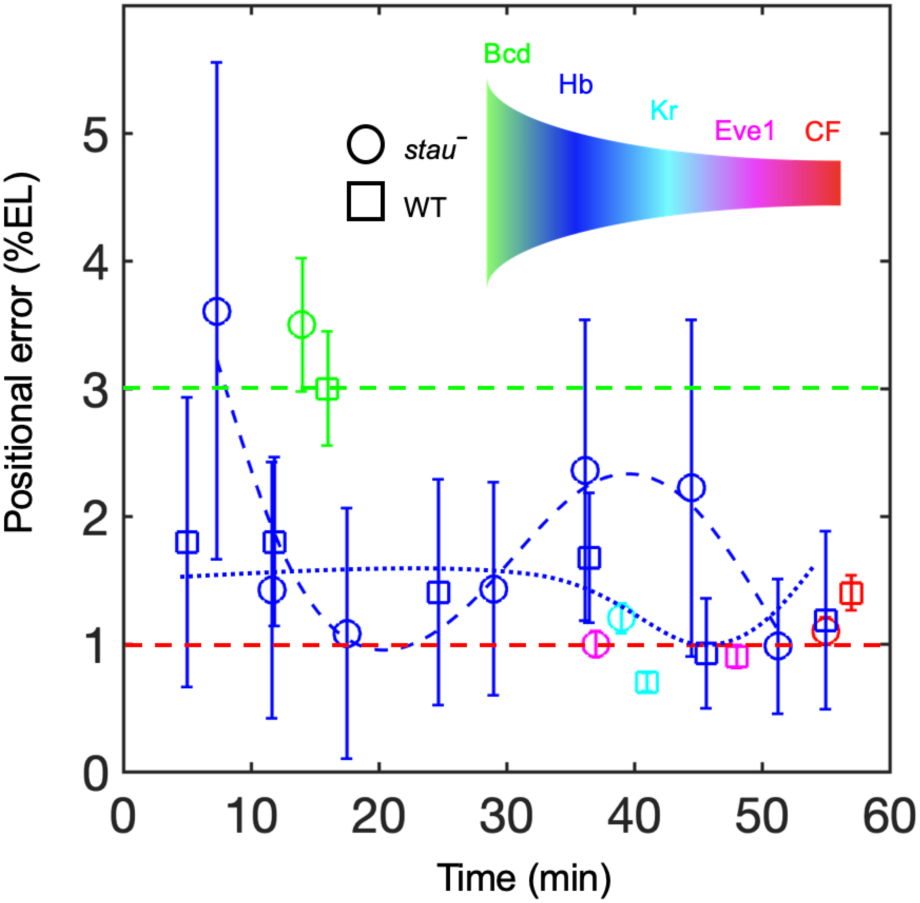
Positional noise filtered from Bcd to CF. Positional errors in Bcd (green), Hb (blue), Kr (cyan), Eve (magenta) and CF (red) of *stau^-^*mutants (circle) and the WT (square) as a function of developmental time in nc14. Green and red dashed lines represent the average positional noise at early and late development times, respectively. Blue lines connect *x_Hb_*in *stau^-^* mutants (dashed line) and the WT (dotted line) to guide the eye.

**Figure 6.**
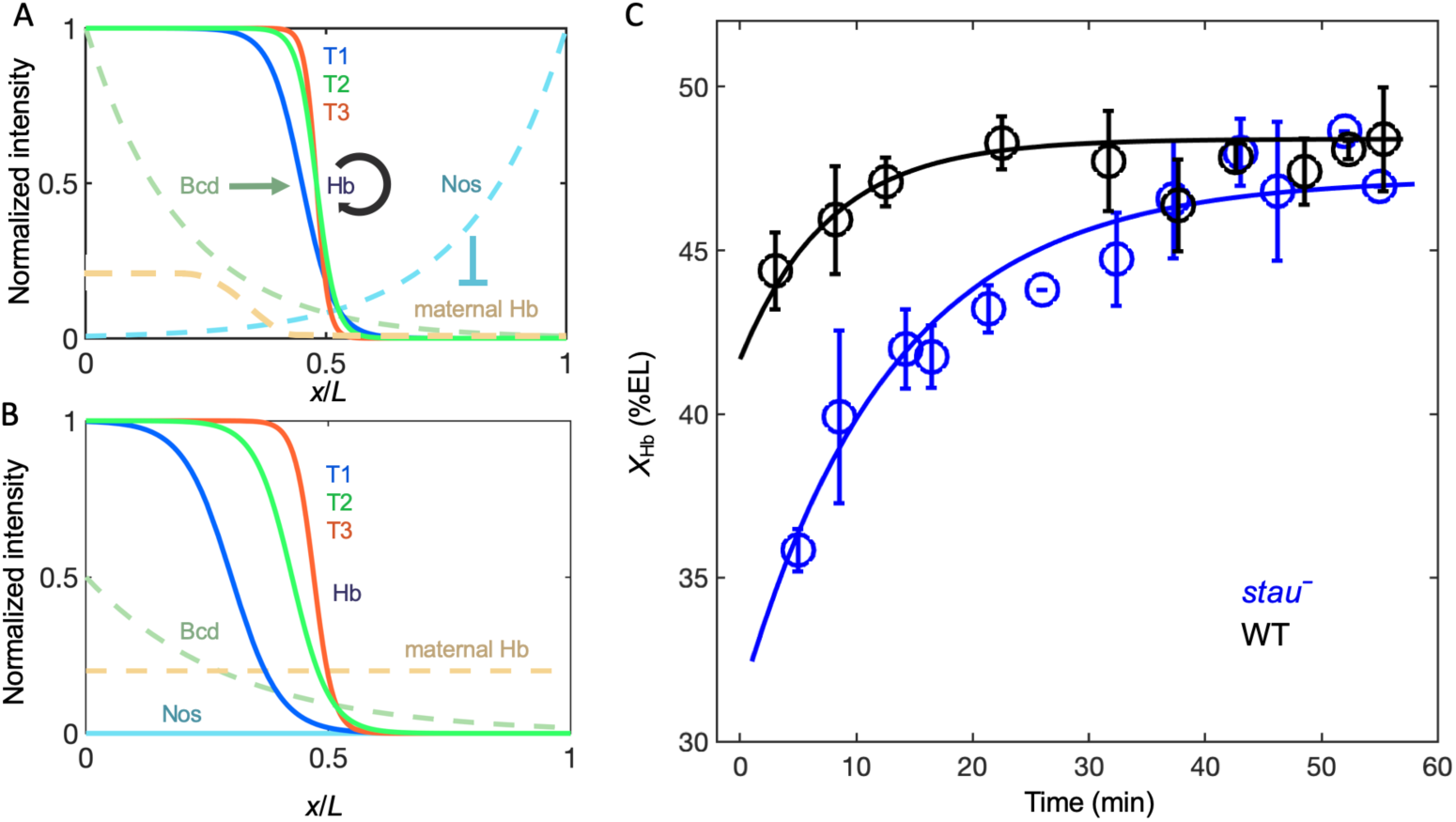
Origin of the large shift in *x_Hb_* in *stau^-^*mutants. (A-B) Schematic of maternal gradients and Hb dynamic expression in the WT (A) and *stau^-^* mutants (B). The Hb boundary shifts posteriorly from early (T1, blue) to middle (T2) and late (T3, red) nc14. It mainly shifts in the first half of hour in nc14 (T1-T2), and the shift amount in *stau^-^* mutants is much greater than in the WT. This difference is probably due to the change in the maternal gradients. Hb is activated by Bcd (dark green) and self-activated by itself. The maternal Hb (orange) is repressed by Nos (azure). In *stau^-^* mutants, the Bcd gradient decreases in amplitude and increases in length. The depletion of the Nos gradient flattens the maternal Hb gradient. (C) The mathematical model fitting (lines) agrees well with the measured shift in *x_Hb_* in *stau^-^* mutants (blue circles) and the WT (black circles). Error bars represent the standard deviation of *x_Hb_* in each time window.

## Materials and Methods

### Fly strains

Bcd-GFP intensity was measured in the fly strain *bcd*-*egfp;+;bcd^E1^ and bcd-egfp;stau^HL54^;bcd^E1^*using live imaging. The expression profiles of Hb, Kr, and Eve in immunostained embryos were measured with *w^1118^* and *stau^-^*(*w; stau^HL54^)* mutants. The *stau^HL54^* mutant was generated from *w^1118^* by a point mutation (replacing the base T in the intron of the fifth exon of *stau* with base) using CRISPR/Cas9 (Bassett and Liu, 2014).

### Immunostaining

All embryos were collected at 25°C, dechorionated, and then heat fixed in 1x TSS (NaCl, Triton X-100). After at least 5 min, the embryos were transferred from the scintillation vial to Eppendorf tubes and vortexed in 1:1 heptane and methanol for 1 min to remove the vitelline membrane. They were then rinsed and stored in methanol at −20°C. The embryos were then stained with primary antibodies, including mouse anti-Hb (Abcam, ab197787), guinea pig anti-Kr, and rat anti-Eve (gifts from John Reinitz). Secondary antibodies were conjugated with Alexa-647 (Invitrogen, A21240), Alexa-488 (Invitrogen, A11006), and Alexa-555 (gift from John Reinitz). To prevent cross-binding between the rat and mouse primary antibodies, the embryos were incubated in guinea pig first and then rat primaries, followed by their respective secondaries; subsequently, the embryos were treated in blocking buffer before the mouse primary and secondary antibodies were applied. Finally, the embryos were stained with DAPI (Invitrogen, D1306).

### 3D imaging and analysis

Embryos stained and washed together in the same tube were mounted in agarose (Invitrogen™ E-Gel™ EX Agarose Gels, 1%). Samples were imaged on a Zeiss Z1 light-sheet microscope. Images were taken with a W Plan Apo 20X/1.0 water immersion objective and with sequential excitation wavelengths of 638, 561,488 and 405 nm. For each embryo, two z-stacks of images (1920×1920 pixels, with 16 bits and a pixel size of 286 nm at 0.8 magnified zoom) with 1 µm spacing were taken from two opposite sides by rotating the embryos 180° (Figure 1-figure supplement 1A). Image analysis routines were implemented in customized MATLAB codes (MATLAB 2018a). The two z-stacks of each embryo acquired from opposite directions were registered with the autocorrelation algorithm (Figure 1-figure supplement 1B), which is based on the correlation coefficient reflecting the similarity between two images. The 3D embryos were reconstructed with these fused z-stack images by the 3D interpolation algorithm. The dorsal and ventral side was determined and ranked by the distance from the surface vertical to the AP axis. 2D images were extracted from the 3D embryos by rotating slices with 5° intervals at different orientations along the AP axis (Figure 1-figure supplement 1C). The gene expression profiles were extracted from these slices.

### Live imaging and analysis

Embryos were collected at room temperature for 1 h, dechorionated in 100% bleach (4% NaClO) for 3 min, glued on the cover glass of a petri dish 30 mm in diameter (NEST, 801002) and covered with halocarbon oil (Halocarbon 700). The Bcd gradient in the midcoronal plane at 16 min into nc14 was measured as reported previously. In brief, embryos were imaged at 22°C on a TPM built in house. The excitation laser was 25 mW in average power and 970 nm in wavelength. The objective was a Zeiss 25X (NA=0.8 in air) oil/water immersion objective. Emission fluorescence was collected with a gallium-arsenide-phosphide (GaAsP) with a quantum yield of more than 40% and dark counts less than 4000/s at 25 °C. For each embryo, three images (512*512 pixels with a pixel size of 460 nm, bit depth of 12 bits, scan speed of 4 ms/line) were taken sequentially along the A-P axis and stitched together. In each session, embryos from the fly strain *bcd*-*egfp;+;bcd^E1^ and bcd-egfp;stau^HL54^;bcd^E1^* were mounted on the same slides and measured side by side.

For the dynamic Bcd gradient measurements, the embryos were collected and imaged in the midsagittal plane at 25 °C on a Nikon A1RSi+ confocal microscope with a Nikon Plan Apo λ 20X/0.75 air objective. The fluorescence was excited at a wavelength of 488 nm and collected with a GaAsP detector. A maximum-intensity-projected z-stack of 5 images (1024×1024 pixels with a pixel size of 620 nm, bit depth of 12 bit, spacing of 1 µm) around the largest plane was acquired at each time point. Each of the images was averaged on two sequential acquisitions. In each session, at most three embryos were picked for imaging to guarantee the time resolution (80 s, scan speed: 2.3 s/frame). The background was the average of the background fluorescence measured with at least ten *w^1118^* embryos under the same conditions. To correct for the potential imaging differences in different sessions, the control samples were prepared with the hand-peeling protocol to preserve the fluorescence of the Bcd-GFP of the collected embryos. Embryos at the interphase of nc13-14 were selected and imaged in advance of each imaging session. Imaging analysis was processed with customized MATLAB codes (MATLAB 2018a).

## Acknowledgements

We thank John Reinitz for antibodies and Lu Wang for assistant in *Drosophila* experiment. This project is supported by the National Natural Science Foundation of China 31670852 and 100-talent plan of Peking University. The *Drosophila* lab used in this project is supported by Peking-Tsinghua Center for Life Sciences.

## Author contributions

Zhe Yang, Investigation, Visualization, Methodology, Data curation; Hongcun Zhu, Investigation, Visualization, Methodology, Data curation; KaKit Kong, Jialong Jiang, Jingchao Zhao, Investigation, Modeling, Software; Jiayi Chen, Xiaxuan Wu, Peiyao Li, Investigation, Data curation; Feng Liu, Conceptualization, Resources, Supervision, Data curation, Software, Funding acquisition, Investigation, Visualization, Methodology, Writing—original draft, Writing—review and editing.

## Supplementary materials

### Model

For simplification, Hb patterning is considered as a one-dimensional reaction-diffusion system assuming the degradation rate and diffusion constant are *β* and *D*, respectively:

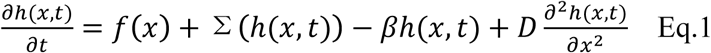

where *h(x, t)* denotes the concentration of Hb at time *t* and the AP axis coordinate *x*, *f(x)* represents the activation of the stable maternal gradient such as Bcd, Σ(h(x,t)) stands for self-activation and takes the form of a step function for simplicity:

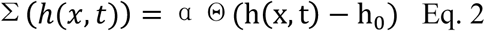

where ϴ is the heaviside function. This indicates that the self-activation remains off until it is fully activated after the Hb concentration exceeds a certain threshold *h_0_*. Eq.1 has a variational form

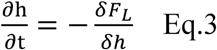

where F_L_ is the Lyapunov function for *h(x,t)*:

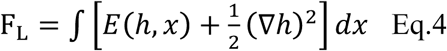

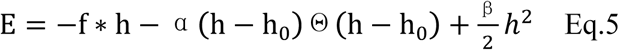

Hence the production of Hb is analogic to phase transition, in which *h* is the order variable, *f* is the external field, and *F_L_* is the free energy. When *f* is in the interval 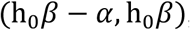, the free energy density *E* has the form of a double well (Fig. S11). The minimal points of the two wells correspond to the solutions 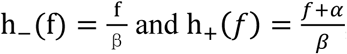, which correspond to the two stable states of the system, i.e., the highest and lowest expression of Hb.

As the system approaches the steady state, the wave front of Eq.1 corresponding to the Hb boundary is generally moving, and its velocity and direction of motion are determined by the free energy density distribution of the system. Let the wave front move at speed *c* and 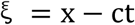, then Eq.1 becomes

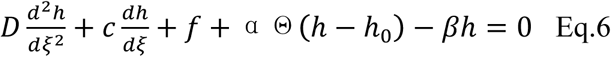

The wave front solution of Eq.6 can be decomposed into two regions. The outer zone is formed in a region far from the wave front, and the system varies little with the independent variable in this area, i.e., Hb concentration is almost flatten in the region away from the boundary. At this time, only the stable solution of the equation is considered.

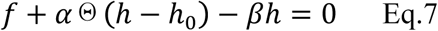

The region near the wave front, i.e., the Hb boundary, constitutes an inner zone where the Hb concentration changes rapidly. We assume the external field *f* is stable: in the wave front propagation region, and the change of the external field *f* is much smaller than the change of Hb: 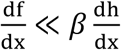. We replace the variable so that *u*=*h-f/β*

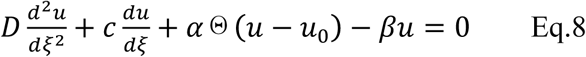

The wave front solution of Eq.6 can be obtained with a standard method(Murry, 2005), and the wave velocity

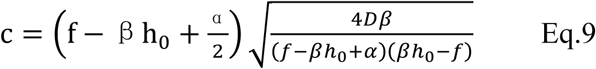

The relationship is as shown in Figure S11B.

Hence at the static wave front, i.e., 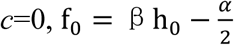 We can define 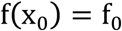 when 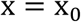. At around 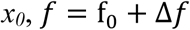, the wave front velocity *c* is approximately linear with the response of the external field 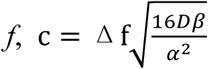.

Expand *f(x)* around *x*_0_

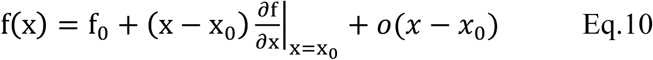

Only consider the first order and let 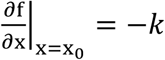, then

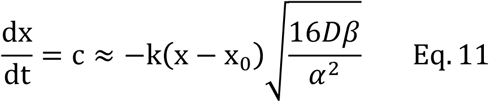

The dynamic change of the boundary position of Hb to reach the stable position *x_0_* follows the following formula:

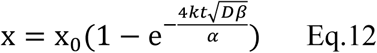

Indeed, this formula fits well with the experimental data of the shift of *x_Hb_* from the initial position to the final position in nc14. And the fitted time constant of the WT is 7.6 min (R^2^=0.55), less than that of the *stau^-^* mutant (12.9 min, R^2^=0.95).

Based on Eq. 12, the time constant:

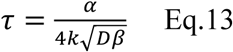

Since the other parameters should be the same for both fly lines, the difference of time constants is attributed to 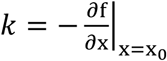. The external field *f* is mainly dependent on the activation of Bcd on Hb, which follows the Hill function with the Hill coefficient of 5(Gregor et al., 2007).

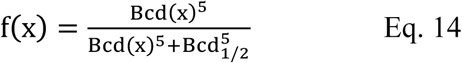

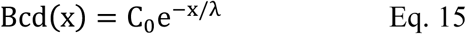

Thus 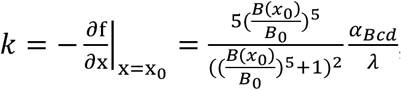, where B(x_0_) and B_0_ are the Bcd concentration at the final boundary position of Hb and the activation threshold of Bcd on Hb, λ and *α_Bcd_* is the length constant of the Bcd gradient and the maximum production rate of Hb activated by Bcd, respectively. After maturation correction on the Bcd gradient, 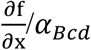 of *stau*^-^ mutants and the WT is showed in Figure S11C. *k_stau_*=0.0082*α_Bcd_* is smaller than *k_WT_=*0.015*α_Bcd_* nM/(min*µm), hence the time constant of *stau*^-^ mutants is larger than that of the WT. Moreover, the ratio of *k_stau_*/ *k_WT_* =0.55 is very close to the ratio of the fitting time constants *τ_WT_*/*τ_Stau_*= 0.59, suggesting this model is consistent with the experiments. Based on previous modeling, the degradation rate *β* =0.1 /min and diffusion constant *D=12* µm^2^/min(Jaeger et al., 2004; Papatsenko and Levine, 2011), plug these values together with the calculated *k* and the fitted time constant into Eq.13, we can obtain the ratio of the maximum production rate 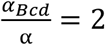 or 2.2 for the WT and the *stau*^-^mutants, respectively.

---

**Table S1.**
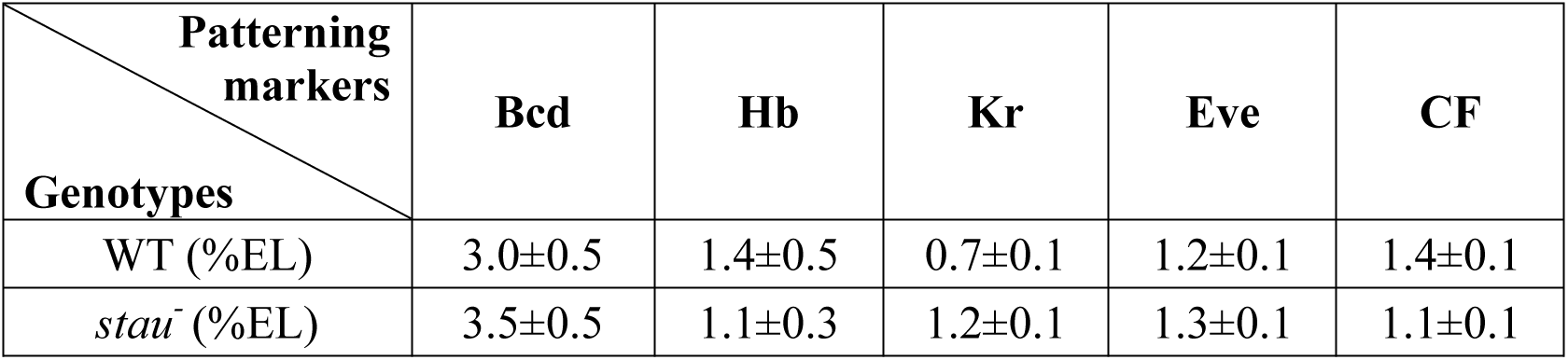
Comparison of the positional noise of patterning markers including average Bcd gradients at 16 min into nc14, posterior boundary of the anterior Hb domain at 12 min into nc14, the anterior boundary of Kr at 40 min into nc14, the 2^nd^ peak of Eve at 42.5 min into nc14, and CF at 56 min into nc14. The standard deviation of the average positional noise is calculated based on bootstrapping.

| **Patterning**<br>**markers**<br>**Genotypes** | **Bcd** | **Hb** | **Kr** | **Eve** | **CF** |
| --- | --- | --- | --- | --- | --- |
| WT (%EL) | 3.0±0.5 | 1.4±0.5 | 0.7±0.1 | 1.2±0.1 | 1.4±0.1 |
| *stau*<sup>-</sup> (%EL) | 3.5±0.5 | 1.1±0.3 | 1.2±0.1 | 1.3±0.1 | 1.1±0.1 |

## Supplementary Figures

**Figure 1-figure supplement 1.**
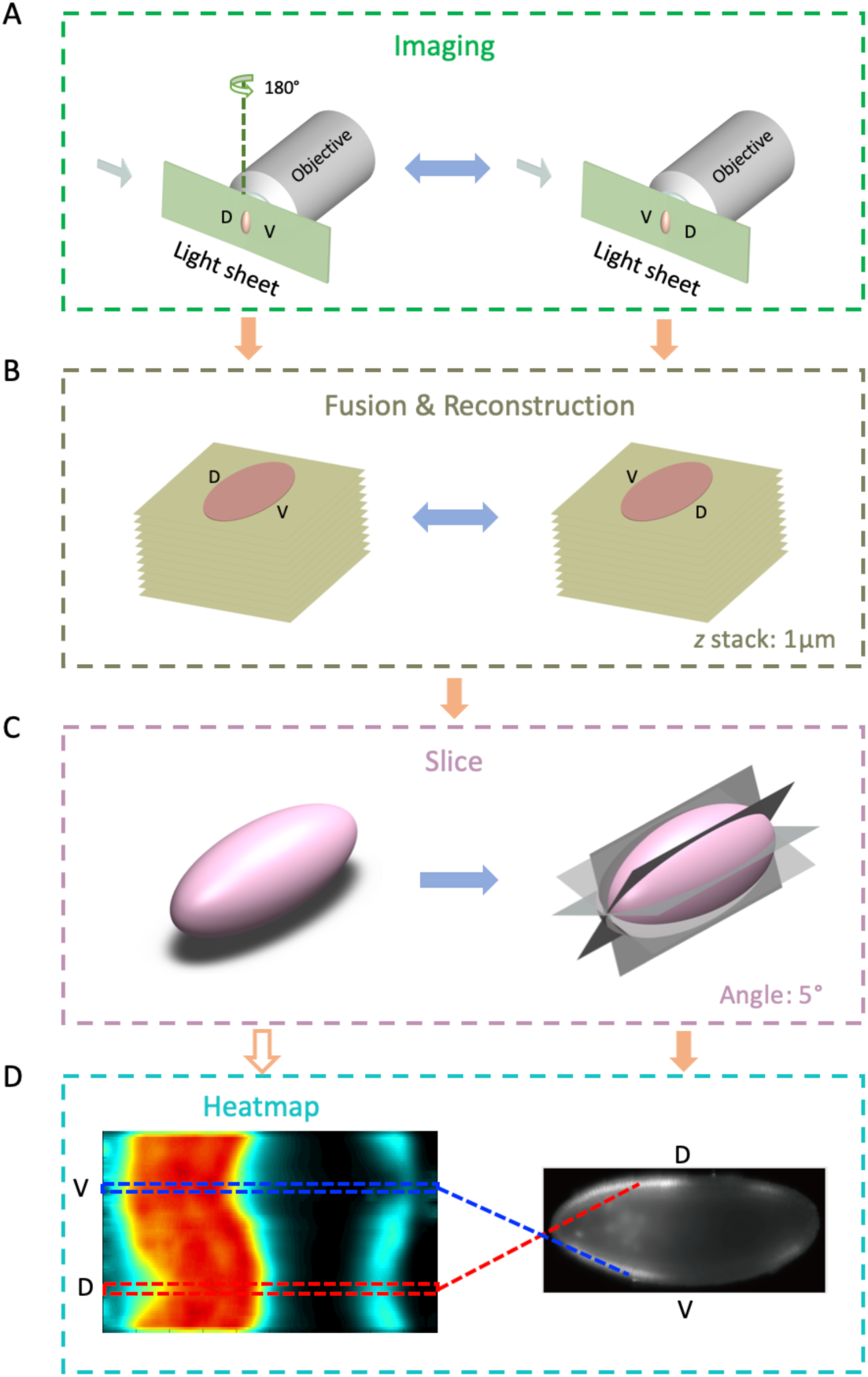
3D imaging measurement procedure. (A) Each embryo was rotated 180° to be imaged from two opposite directions with a light-sheet microscope. (B) The resultant z-stacks were corrected for intensity decay as a function of the depth then fused together to reconstruct a 3D embryo. (C) The expression profiles were extracted from the 2D plane rotating along the A-P axis of the embryo. (D) These profiles were normalized with the maximum intensity of the whole embryo and assembled to form the final heat map (left). The dorsal and ventral profiles are extracted from a representative lateral view of the embryo image (right) sliced from the reconstructed 3D embryo.

**Figure 1-figure supplement 2.**
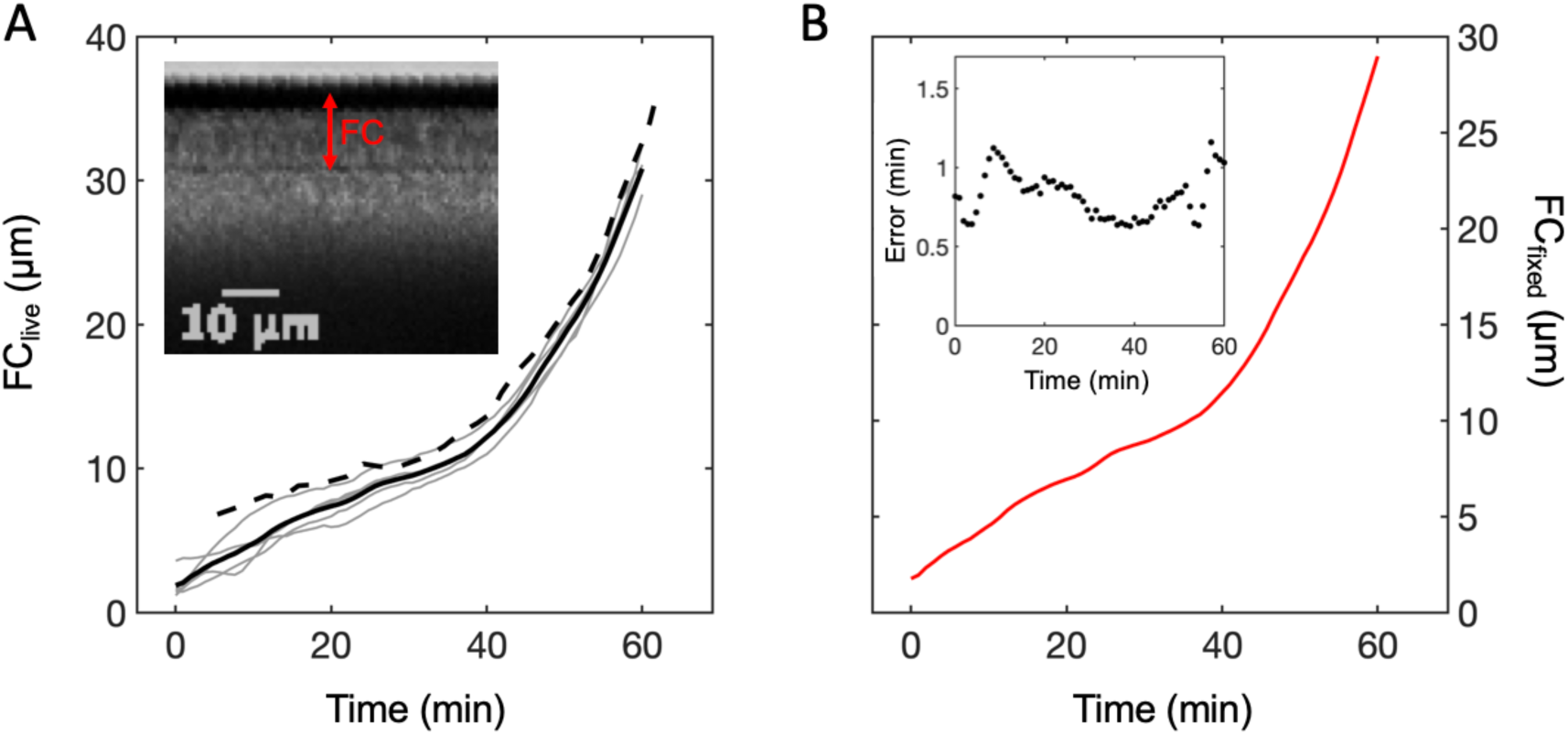
Age determination of fixed *Drosophila* embryos. (A) The depth of the membrane furrow canal (FC) on the dorsal side of the w1118 embryos measured with live imaging as a function of the developmental time into nc14 (*t*). Each gray line represents the curve of FC vs. *t* measured with one embryo. The average curve of FC vs. *t* of 5 *w^1118^* embryos (black line) is slightly different from that of Oregon-R (Dubuis et al., 2013) (Dashed line). Inset shows a middle dorsal part of a representative image of live *w^1118^* embryo and the corresponding FC. (B) The calculated depth of the membrane furrow canal (FC) on the dorsal side of the fixed *w^1118^* embryos with a shrinkage ratio of 94% as a function of the developmental time into nc14 (*t*). Inset shows the error in determining the embryo age of fixed embryos with the measured curve of FC vs. *t*.

**Figure 1-figure supplement 3.**
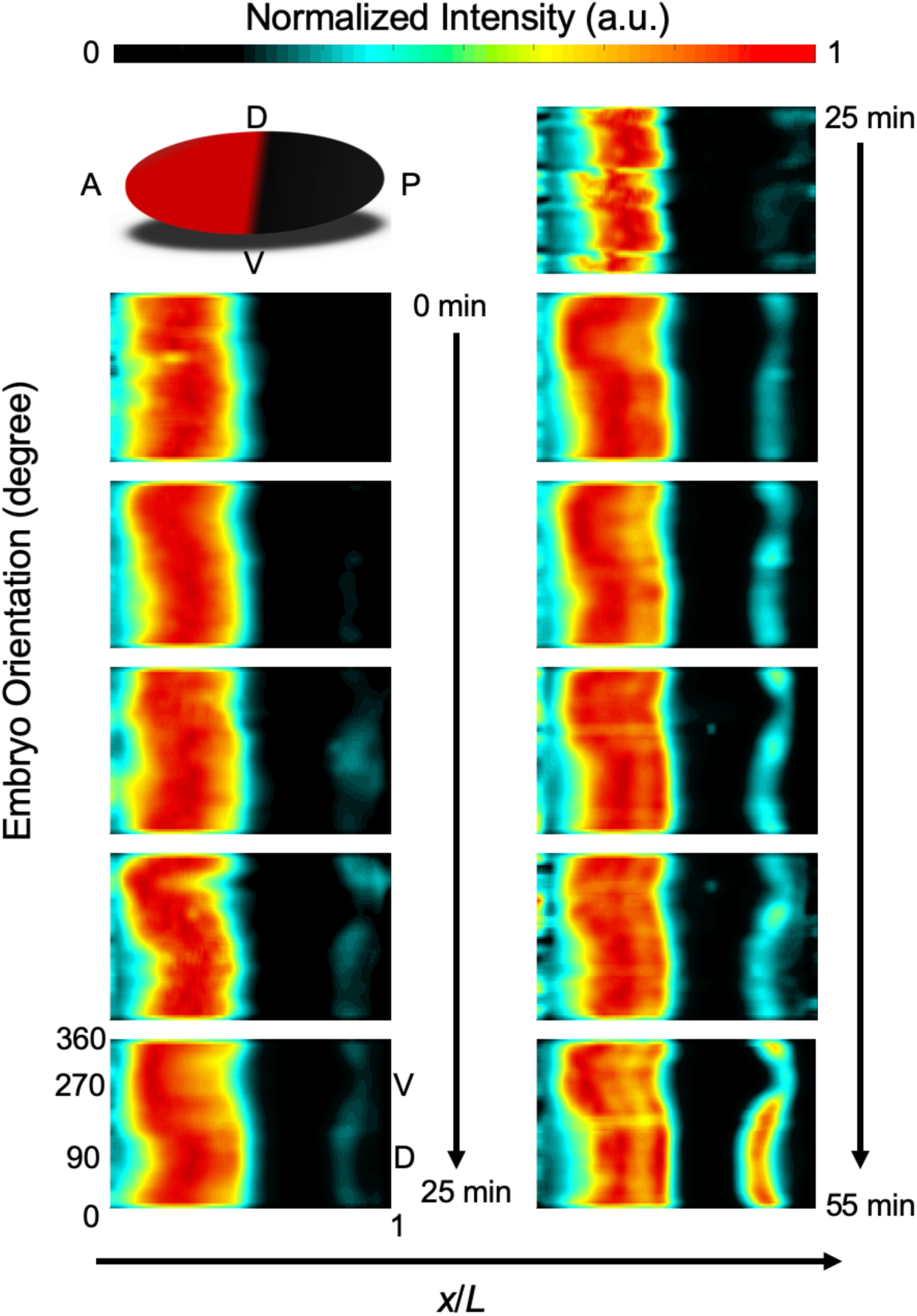
Heat map of the Hb profiles of the WT taken from the projection.

**Figure 1-figure supplement 4.**
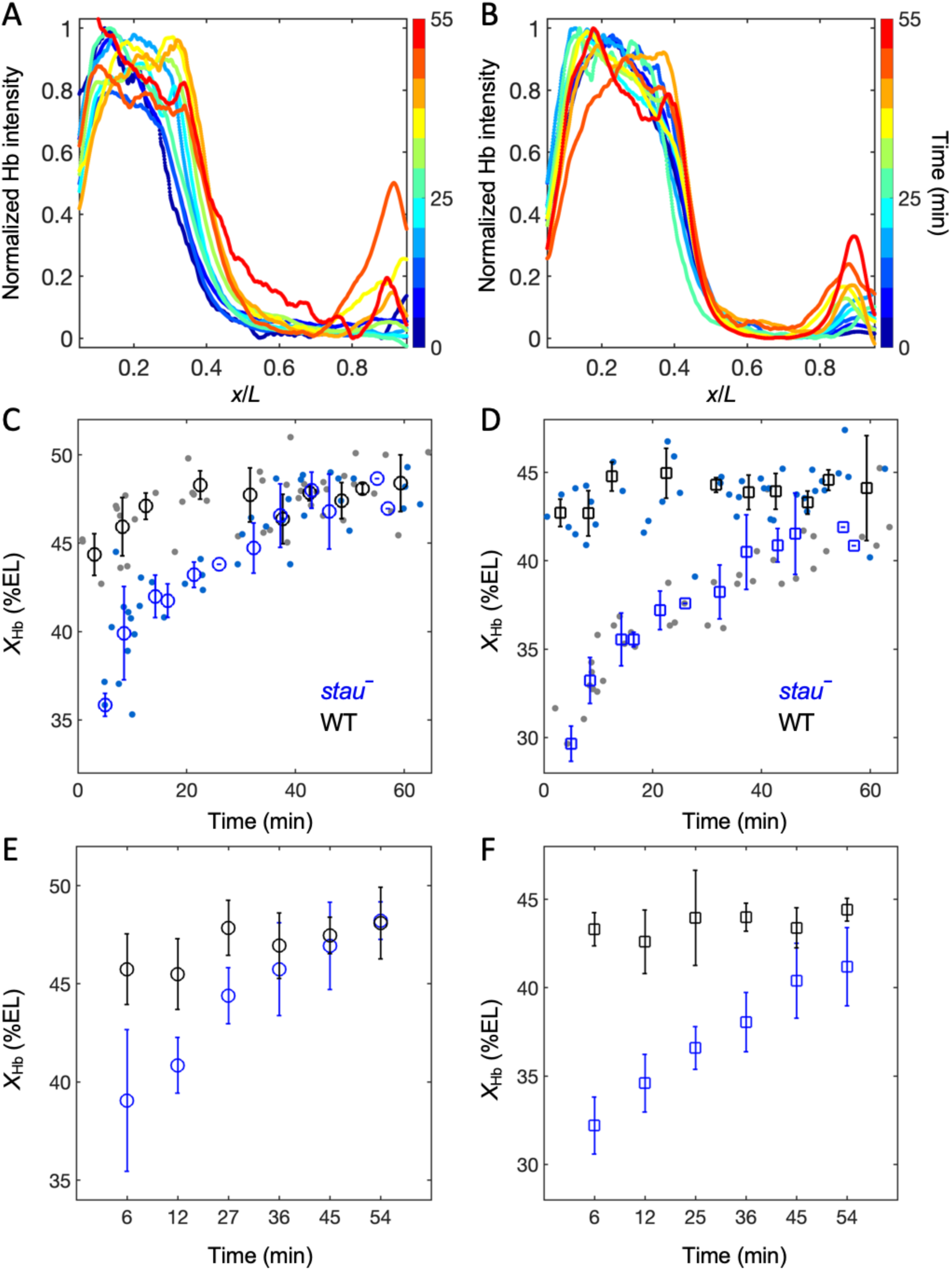
(A-B) The dynamics of the average ventral Hb profiles extracted from 3D imaging on the embryos of the WT (A) and *stau^-^*mutants (B). (C-D) Average dorsal (C) and ventral (D) Hb boundary position (*x_Hb_*) in each 5-min bin of *stau^-^* mutants (blue circle) and the WT (black circle) as a function of the developmental time in nc14, error bars denote the standard deviation of the bin; each dot represents the boundary position of a single embryo. (E-F) Average dorsal (E) and ventral (F) Hb boundary position (*x_Hb_*) in each bin with the same sample size (*N*=6) of *stau^-^* mutants (blue circle) and the WT (black circle) as a function of the developmental time in nc14, error bars denote the standard deviation of the bin.

**Figure 1-figure supplement 5.**
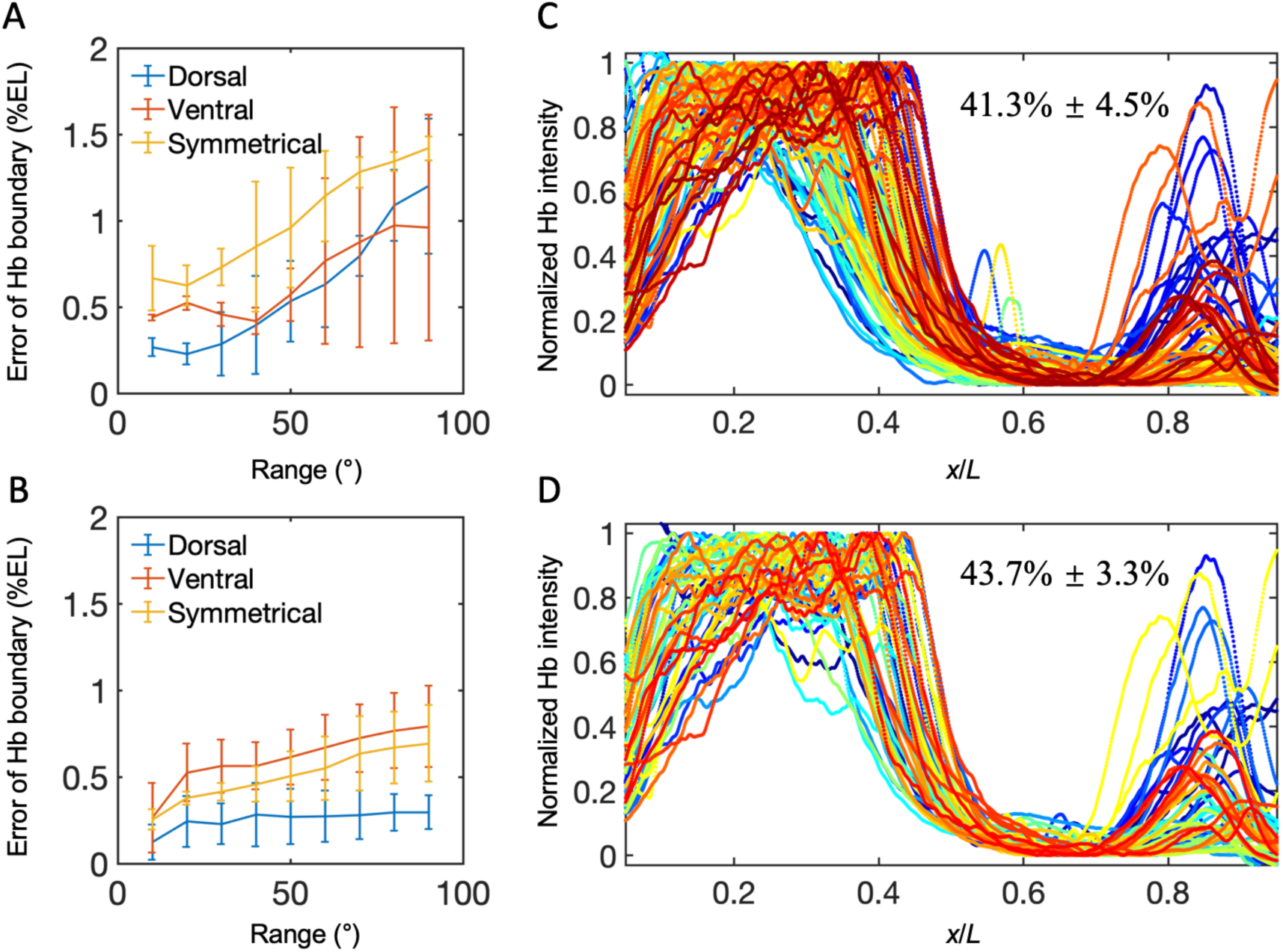
(A-B) The variability of *x_Hb_*extracte*d* from the dorsal, ventral and symmetric profiles as a function of the embryo orientation along the AP axis at 16 min into nc14 (A) and 40 min into nc14 (B). (C-D) All the Hb dorsal profiles without any control on embryo ages and orientations (C) and with the embryo age of 30-50 min and spatial orientation errors less than 30° (D).

**Figure 2-figure supplement 1.**
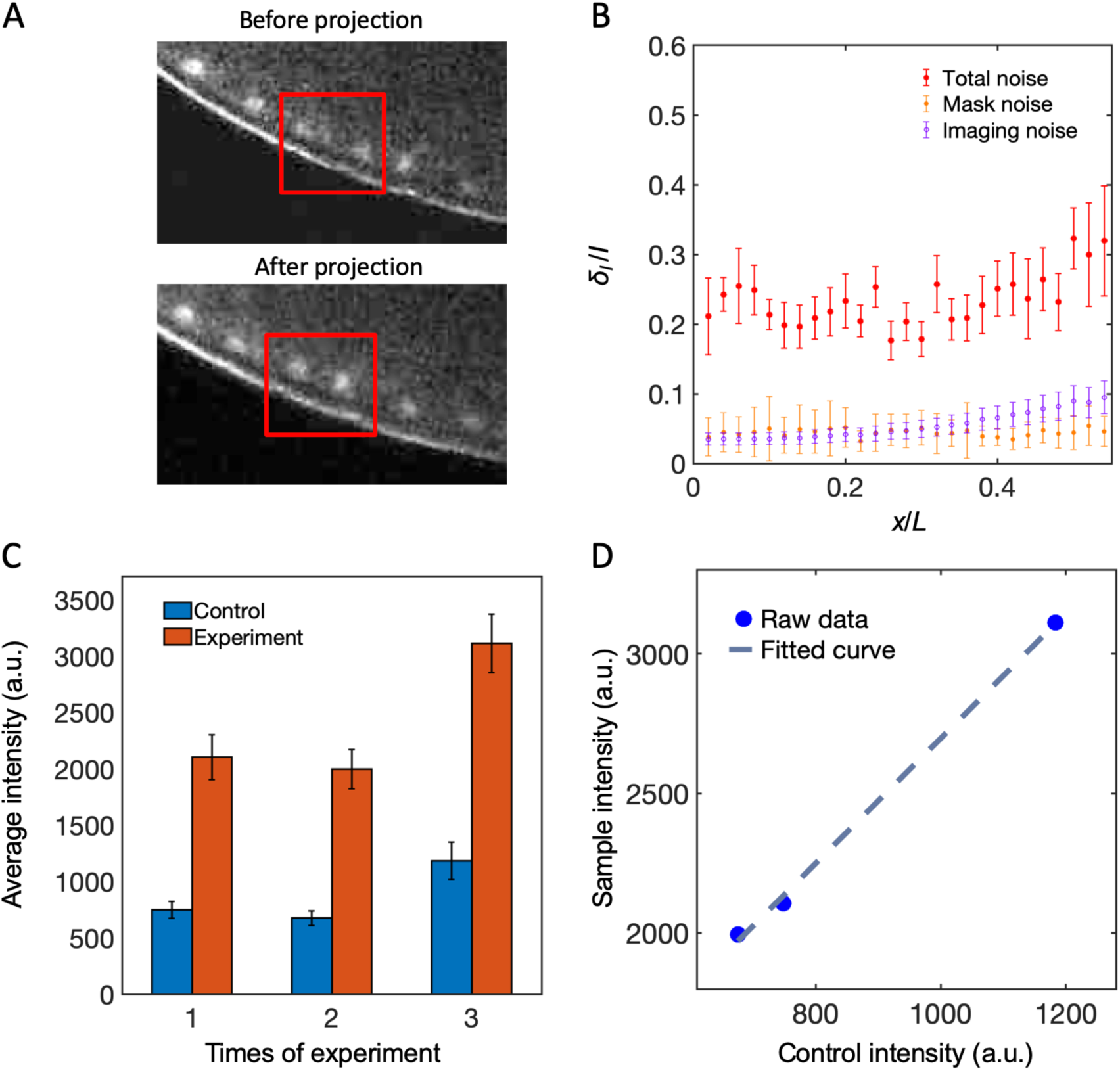
(A) Comparison of the maximum projected image and the single image in the mid-sagittal plane from a z-stack of images of the fly embryos expressing Bcd-GFP in nc12. (B) The nuclear mask noise from imaging segmentation and the imaging shot noise is much smaller than the total Bcd-GFP gradient noise. (C) Comparison of the average nuclear fluorescence intensity of the control sample and the embryos measured in three independent sessions. (D) The average nuclear fluorescence intensity of the embryos is linear corrected with that of the control sample in three measurement sessions. Blue circle represent data from each session, and dash line represent the linear fitting.

**Figure 3-figure supplement 1.**
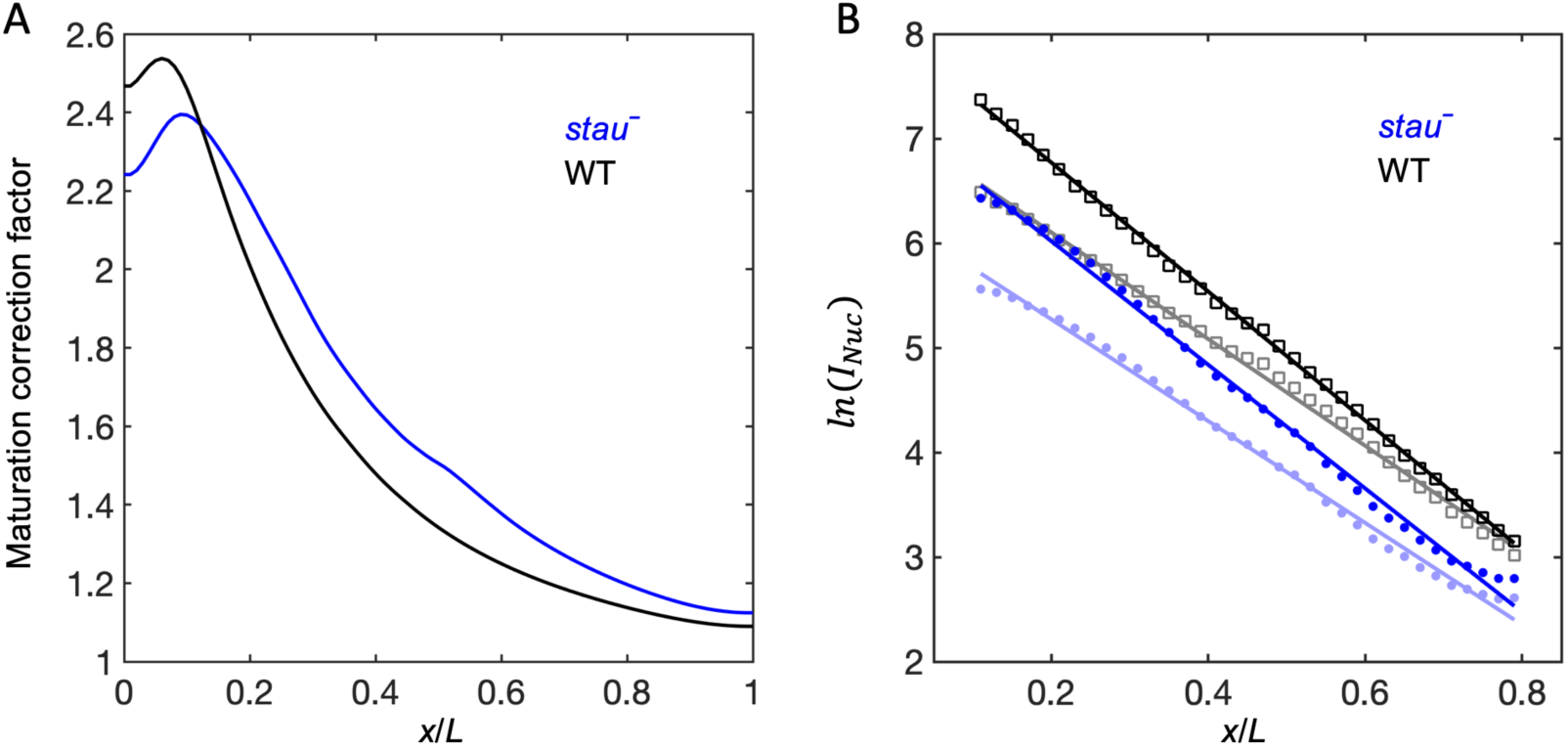
(A) Maturation correction curves of the Bcd gradient. (B) The logarithm of the intensity as a function of the fractional embryo length and the linear fit with before maturation correction (light color) and after maturation correction (dark color).

**Figure 4-figure supplement 1.**
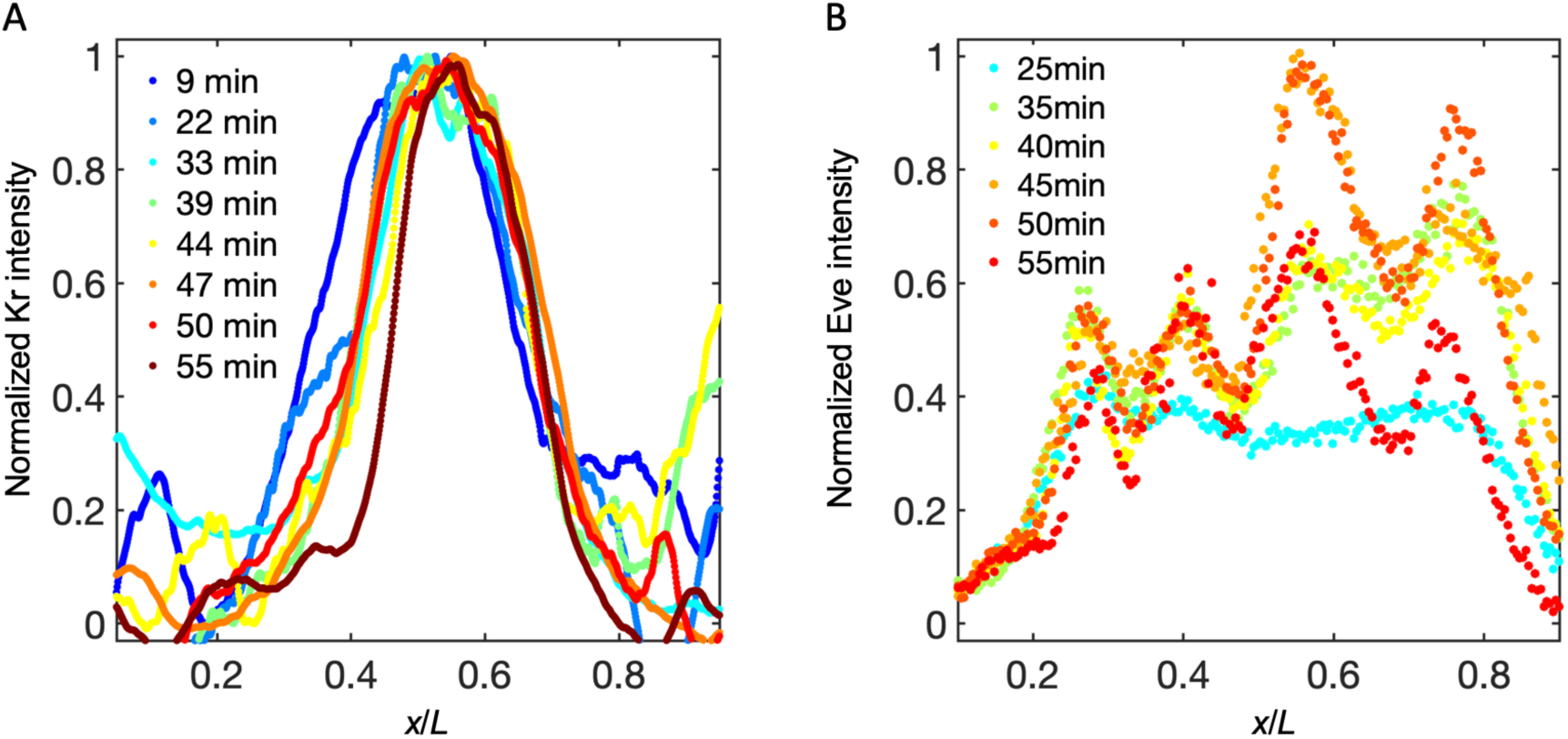
Measurement results of Kr and Eve in *stau^-^* mutants. (A-B) Dynamics of the Kr (A) and Eve (B) profiles in nc14.

**Figure 4-figure supplement 2.**
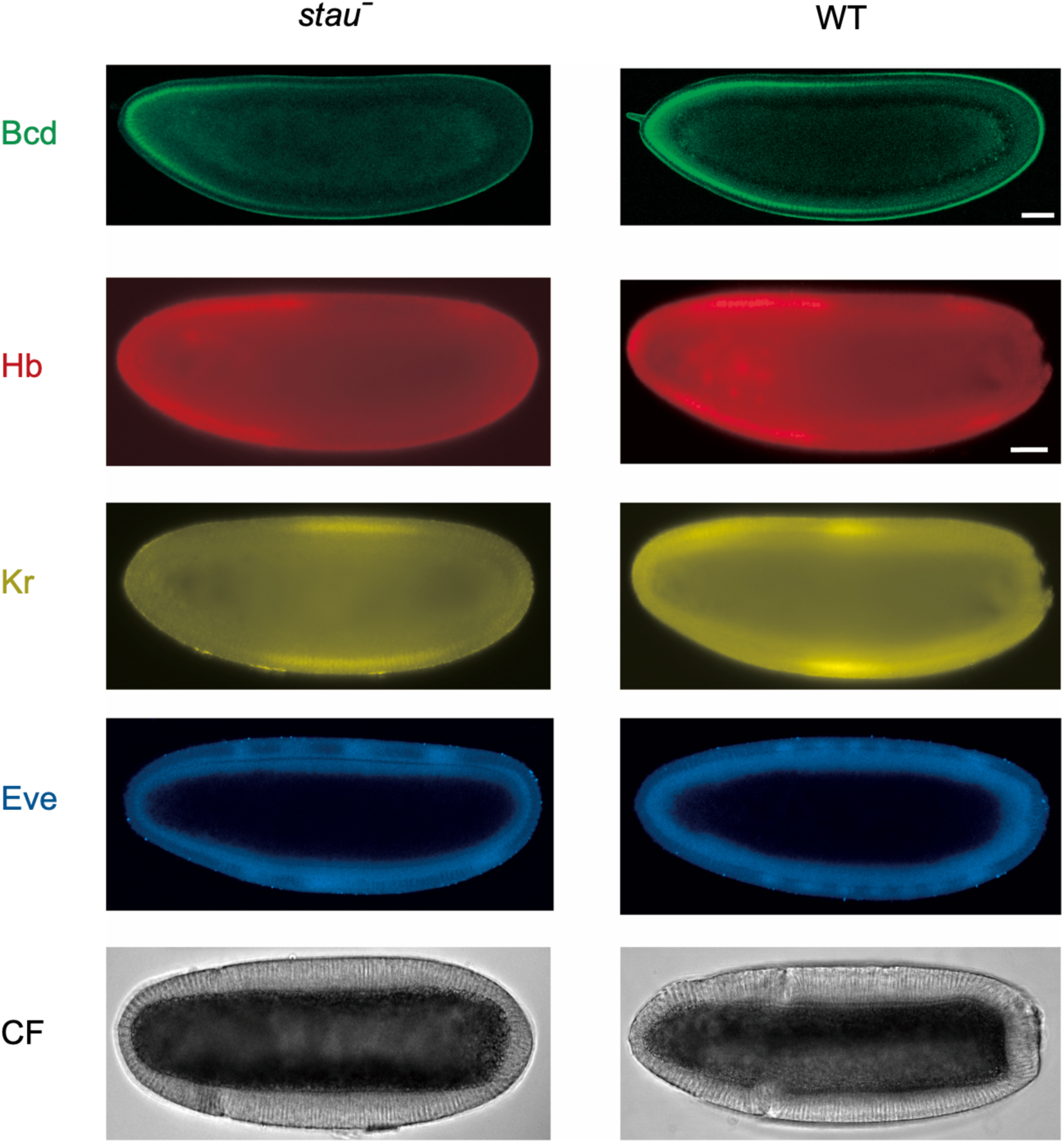
Representative raw images of Bcd-GFP (green), Hb (red), Kr (yellow), Eve (blue), and CF (black) of *stau^-^*mutants (left) and the WT (right). The top two figures are living images of Bcd-GFP at 16 min into nc 14, and the next three rows of figures are the immunofluorescence images of Hb, Kr, and Eve at around 35 min into nc 14. The last row is the bright field image of CF at around 56 min into nc 14. The scale bars represent 40 µm. The first one is for the top two images and the second one for the rest. The second one is slightly longer due to the shrinkage after heat fixation.

**Figure 5-figure supplement 1.**
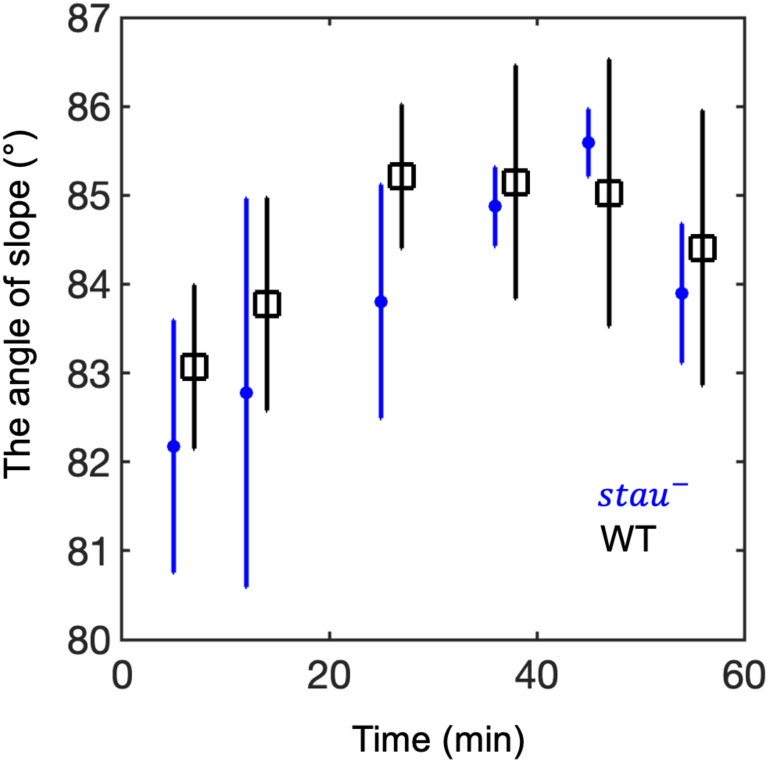
The average Hb boundary slopes of the *stau^-^* mutants (blue circle) and the WT (black circle) as a function of the developmental time in nc14, error bars denote the standard deviation of each bin with the same sample size (*N* = 6). The time points of the WT have a 1-min offset in the time axis for visual display.

**Figure 6-figure supplement 1.**
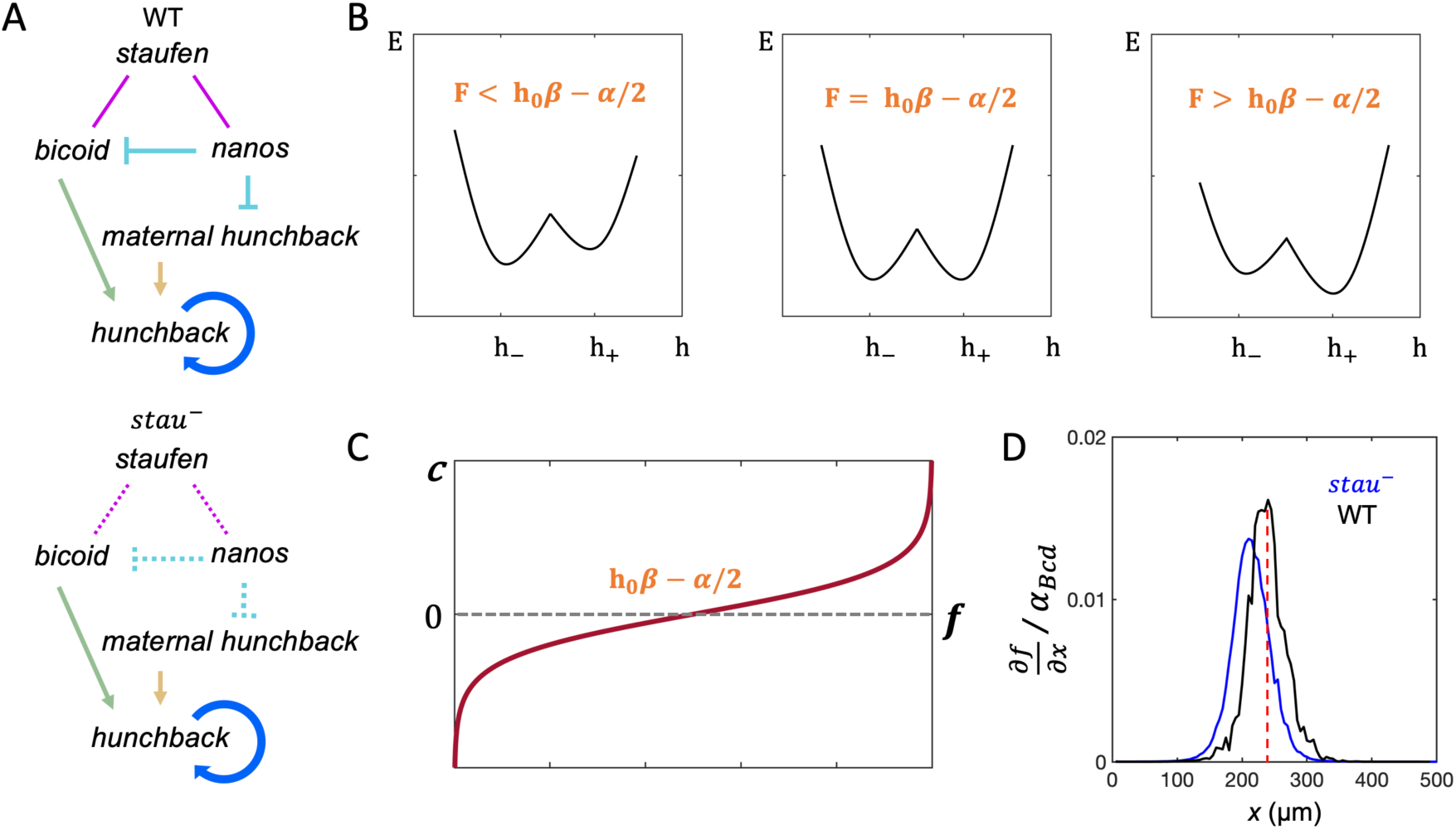
Mathematical modeling of the shift of *x_Hb_*. (A) The regulation network of maternal genes and Hb in the WT (top) and the *stau^-^* mutants (below). The arrow (->) and the symbol (**—|**) represent the activation and repression, respectively. The dashed line represents the missing regulatory interactions. (B) Free energy density *E* as a function of the Hb concentration *h* when the free energy is tuned. *h*_-_ and *h*_+_ represent two stable solutions. (C) The wave front velocity *c* as a function of external field *f.* (D) The calculated 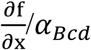 of *stau*^-^ mutants (blue) and the WT (black) as a function of *x* (coordinate along the AP axis with the anterior pole as 0). The red dashed line is set approximately at the stable position *x*_0_ of the WT and *stau^-^* mutants as the two positions are very close to each other.

Figure 1-video 1. The z-stack of the immunofluorescence images of DAPI of the *stau^-^*mutant embryo after fusion in 3D imaging.

Figure 1-video 2. The z-stack of the immunofluorescence images of Hb of the *stau^-^*mutant embryo after fusion in 3D imaging.

